# Systems-wide analysis of the ROK-family regulatory gene *rokL6* and its role in the control of glucosamine toxicity in *Streptomyces coelicolor*

**DOI:** 10.1101/2023.09.20.558697

**Authors:** Chao Li, Mia Urem, Chao Du, Le Zhang, Gilles P. van Wezel

## Abstract

Streptomycetes are saprophytic bacteria that grow on complex polysaccharides, such as cellulose, starch, chitin and chitosan. For the monomeric building blocks glucose, maltose and *N*-acetylglucosamine (GlcNAc), the metabolic pathways are well documented, but that of glucosamine (GlcN) is largely unknown. *Streptomyces nagB* mutants, which lack glucosamine-6-phosphate deaminase activity, fail to grow in the presence of high concentrations of GlcN. Here we report that mutations in the gene for the ROK-family transcriptional regulator RokL6 relieve the toxicity of GlcN in *nagB* mutants, as a result of elevated expression of the Major Facilitator Superfamily (MFS) exporter SCO1448. Systems-wide analysis using RNA sequencing, ChIP-Seq, EMSAs, 5’RACE, bioinformatics and genetics revealed that RokL6 is an autoregulator that represses transcription of sco1448 by binding to overlapping promoters in the *rokL6*-sco1448 intergenic region. RokL6-independent expression of sco1448 fully relieved toxicity of GlcN to *nagB* mutants. Taken together, our data show a novel system of RokL6 as a regulator that controls the expression of the MFS transporter SCO1448, which in turn protects cells against GlcN toxicity, most likely by exporting toxic metabolites out of the cell.

**IMPORTANCE:** Central metabolism plays a key role in the control of growth and antibiotic production in streptomycetes. Specifically, aminosugars act as signaling molecules that affect development and antibiotic production, via metabolic interference with the global repressor DasR. While aminosugar metabolism directly connects to other major metabolic routes such as glycolysis and cell wall synthesis, several important aspects of their metabolism are yet unresolved. Accumulation of *N*-acetylglucosamine 6-phosphate (GlcNAc-6P) or glucosamine 6-phosphate (GlcN-6P) is lethal to many bacteria, a yet unresolved phenomenon referred to as “aminosugar sensitivity”. We made use of this concept by selecting for suppressors in genes related to GlcN toxicity in *nagB* mutants, which showed that the gene pair of *rok*-family regulatory gene *rokL6* and MFS transporter gene sco1448 forms a cryptic rescue mechanism. Inactivation of *rokL6* resulted in the expression of sco1448, which then prevents toxicity of amino sugar-derived metabolites in *Streptomyces*. The systems biology of RokL6 and its transcriptional control of sco1448 sheds new light on aminosugar metabolism in streptomycetes and on the response of bacteria to aminosugar toxicity.

## INTRODUCTION

Streptomycetes are Gram-positive bacteria with a mycelial lifestyle that reproduce via sporulation. Their large GC-rich genomes encode a plethora of specialized metabolites such as antibiotics, anticancer drugs and many other industrially and medically relevant compounds (1–3). The production of these molecules is closely related to the transition from vegetative to aerial growth during development of the colonies (4, 5). Streptomycetes grow by tip extension and branching of vegetative hyphae, which are divided into multigenomic compartments. Under adverse conditions such as nutrient depletion, a complex developmental program is initiated that results in the formation of an aerial mycelium, and eventually the aerial hyphae differentiate to form chains of unigenomic spores (3, 6). At the onset of development, the vegetative mycelium is partially degraded via programmed cell death (PCD) so as to provide the building blocks necessary for aerial growth in an otherwise nutrient-depleted environment (7, 8).

During cell wall recycling, the constituents of the peptidoglycan (PG), namely *N*- acetylglucosamine (GlcNAc) and *N*-acetylmuramic acid (MurNAc) that make up the PG strands and the cross-linking amino acids, are re-imported into the cell. The lactyl ether substituent of the intracellular MurNAc 6-phosphate (MurNAc-6P) is cleaved by MurNAc-6P etherase (MurQ), yielding *N*-acetylglucosamine 6-phosphate (GlcNAc-6P) (9). GlcNAc is thereby internalized by the phosphoenolpyruvate-dependent phosphotransferase system (PTS) which simultaneously phosphorylates GlcNAc to GlcNAc-6P (10, 11). The next step is GlcNAc-6P deacetylation by *N*-acetylglucosamine 6-phosphate deacetylase NagA forming glucosamine-6-phosphate (GlcN-6P), which is a central molecule at the intersection of multiple metabolic pathways, including glycolysis via conversion to fructose-6-phosphate (Fru-6P) by glucosamine-6-phosphate deaminase NagB (12). GlcNAc plays a critical role in signaling during the control of development and antibiotic production in streptomycetes (13–15). DasR controls aminosugar transport and metabolism and is also a highly pleiotropic repressor of antibiotic production in *Streptomyces* (16–18). Metabolic control of DasR is a key step in the early activation of development and specialized metabolism under nutrient-deprived conditions.

While the metabolic pathway of GlcNAc has been well characterized, little is known of glucosamine (GlcN) metabolism in streptomycetes. The hydrolysis of chitosan and the metabolism of chitosan-derived oligomers (GlcN)_2-3_ are under the control of CsnR (19, 20), a repressor from the ROK (Repressors, ORFs and Kinases) family of transcriptional regulators (21). GlcN oligomers are imported via the ABC-transporter complex CsnEFG-MsiK and are then presumably hydrolyzed and phosphorylated by CsnH, a sugar hydrolase, and CsnK, a ROK-family kinase, respectively (20).

High concentrations of either GlcN or GlcNAc are toxic to *S. coelicolor* in the absence of the enzyme glucosamine-6-phosphate deaminase (NagB) (12). Similar aminosugar sensitivity was also observed for *Escherichia coli* (22, 23). Despite extensive studies focusing on the metabolism of aminosugars, the underlying cause of aminosugar toxicity remains unresolved. Interestingly, when grown in the presence of high concentrations of either GlcN or GlcNAc, *S. coelicolor* Δ*nagB* strains sustain spontaneous second-site mutations that allow the colonies to survive. We previously exploited this principle to identify novel genes related to GlcN transport and metabolism (12, 24). Spontaneous mutations or deletion of *nagA*, preventing the conversion from GlcNAc-6P to GlcN-6P, relieves the toxicity of both GlcNAc and GlcN to *S. coelicolor nagB* mutants (24). Since NagA is not known to play a role in GlcN metabolism, this points at a caveat in our understanding of aminosugar metabolism. In addition to suppressor mutations in *nagA*, two independent suppressor mutations were found in the gene for ROK-family regulator SCO1447 (RokL6), which relieved toxicity specifically of GlcN but not of GlcNAc. ROK (Repressor, ORF and Kinase) family of proteins often play important roles in the (control of) sugar utilization in bacteria, and these regulators are widespread in streptomycetes (21, 25, 26).

Here, we report on the function of RokL6 by a systems-wide approach, using mutational and transcriptional analysis, and *in vitro* and *in vivo* DNA binding studies. This revealed that the gene product of *rokL6* is a ROK-family regulator that specifically represses the adjacent gene sco1448. This gene encodes a putative Major Facilitator Superfamily (MFS) transporter, and we propose that it acts as a pump for toxic substances created during challenge of *nagB* mutants with higher concentrations of GlcN.

## MATERIALS AND METHODS

### Strains, plasmid, and growth conditions

The bacterial strains and plasmids used or constructed in this study are summarized and listed in Table 1, and all oligonucleotides are described in Table 2. *Escherichia coli* was grown and transformed according to the standard procedures (27), with *E. coli* JM109 serving as the host for routine cloning, and *E. coli* ET12567 (28) used for the isolation of non-methylated DNA for transformation into *Streptomyces*. For protein heterologous expression, *E. coli* BL21(DE3) from Novagen was used. *E. coli* was grown in Luria-Bertani (LB) media in the presence of selective antibiotics as required, with the following final concentrations: ampicillin (100 μg/mL), apramycin (50 μg/mL), kanamycin (50 μg/mL) and chloramphenicol (25 μg/mL). *Streptomyces coelicolor* M145 (29) was obtained from the John Innes Centre strain collection and was the parent of all mutants. *S. coelicolor nagB* mutant Δ*nagB* (24) and GlcN-derived *nagB* suppressor mutant SMG1 (30), have been described previously. All *Streptomyces* media and routine techniques are based on the *Streptomyces* manual (31). Phenotypic characterization of *Streptomyces* mutants was carried out on minimal medium (MM) agar plates with different carbon sources as indicated. Soy flour mannitol (SFM) agar plates were used for preparation of spore suspensions and *E. coli* to *S. coelicolor* conjugations. A mixture of 1:1 yeast-extract malt extract (YEME) and tryptic soy broth (TSBT) liquid media was used to cultivate mycelia for protoplast preparation and genome DNA isolation. Growth curves of *Streptomyces* were performed in liquid cultures containing minimal medium (NMMP) (31) with 1% (w/v) glucose or GlcN as the sole carbon source, and the dry weights of the mycelia were measured at different time points.

**Table 1.**
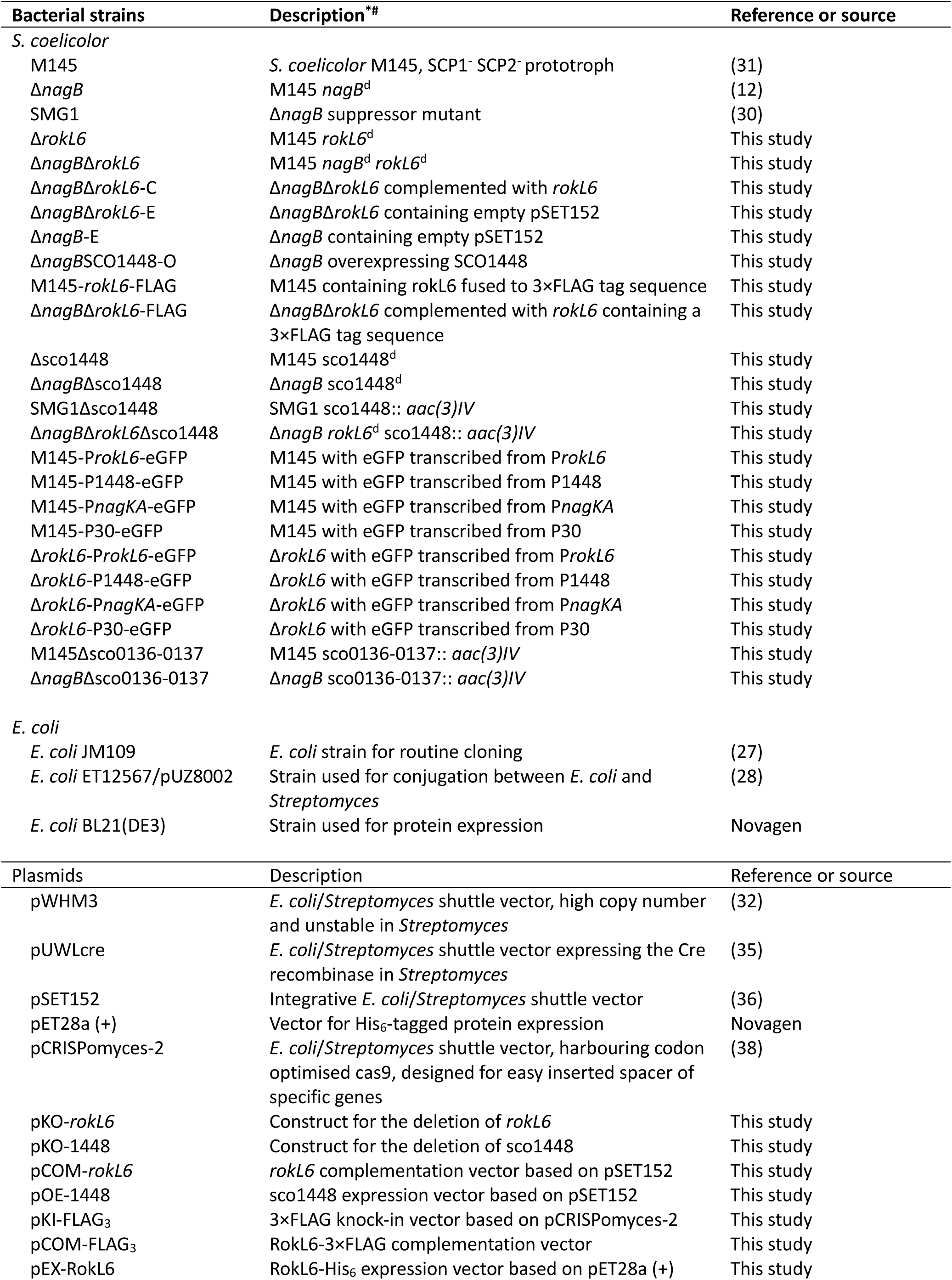
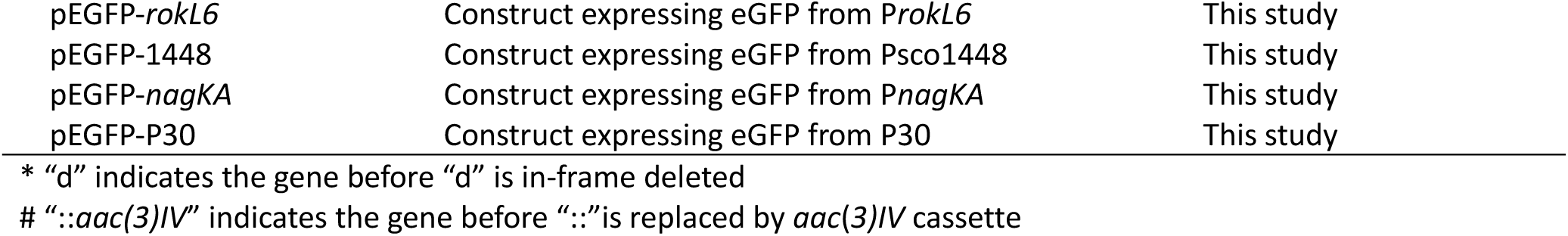
Bacterial strains and plasmids used in this study.

**Table 2.**
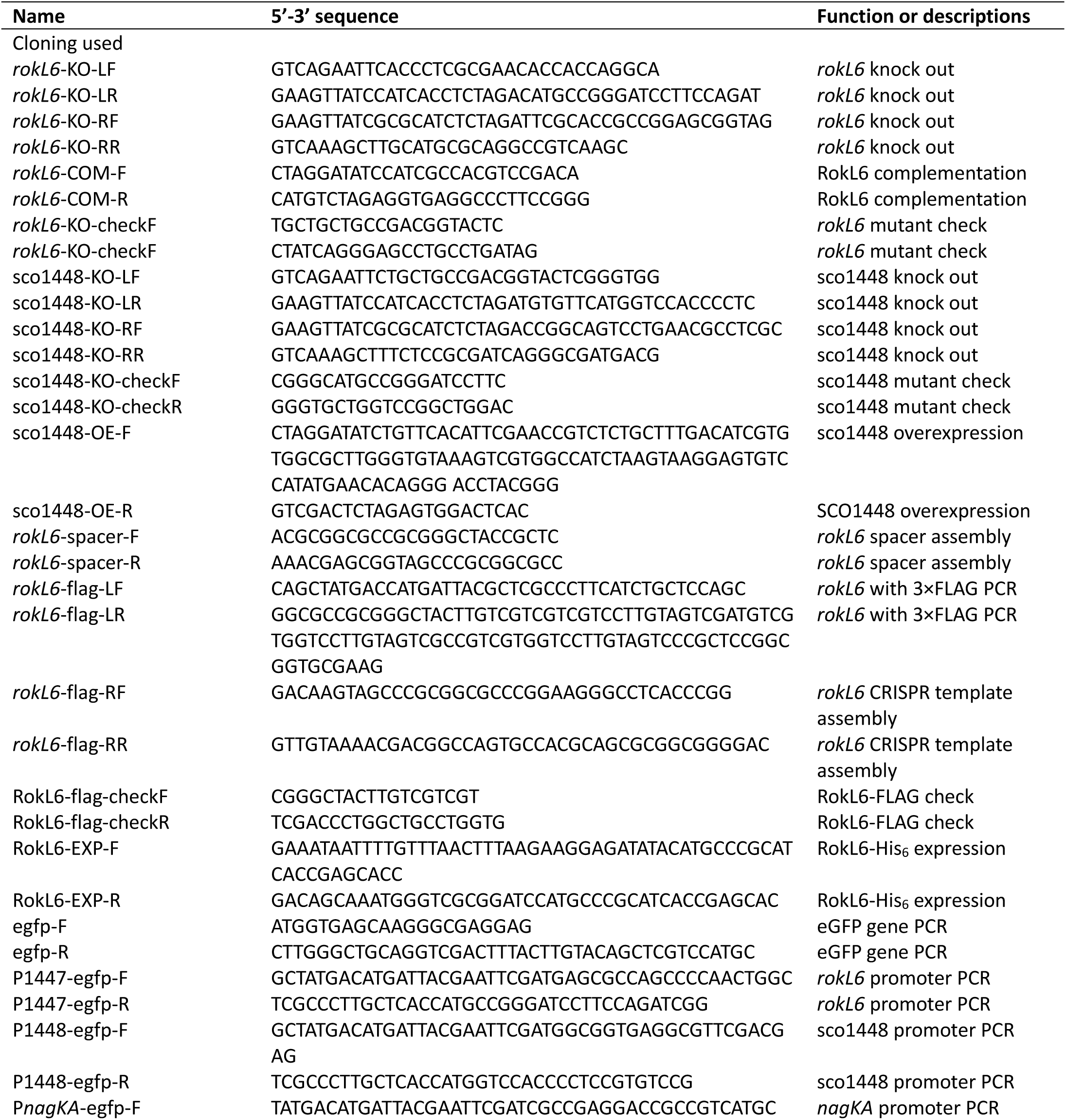
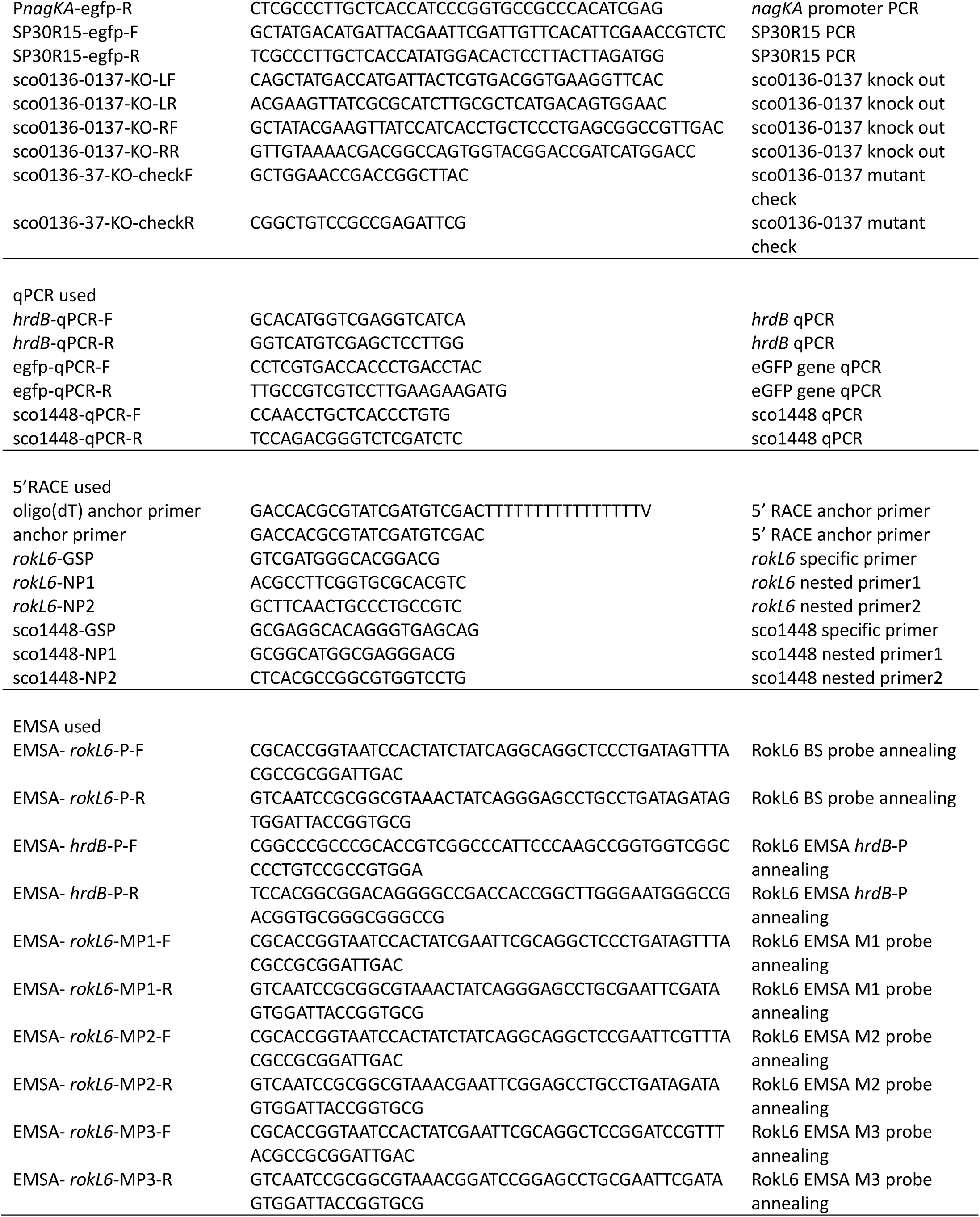
Oligonucleotides used in this study.

### Gene knock-out, complementation and overexpression

As basis for the creation of gene deletion mutants we used the unstable multi-copy plasmid pWHM3 (32, 33), as described previously (12). For the *rokL6* knock-out construct, a 1254-bp 5’ flanking region and a 1366-bp 3’ flanking region were amplified by PCR from the *S. coelicolor* M145 genome, using the primer pairs described in Table 2. The upstream region was cloned as an *Eco*RI-*Xba*I fragment, and the downstream region as an *Xba*I-*Bam*HI fragment, and these two fragments were ligated into pWHM3 from the *Eco*RI and *Bam*HI sites. The apramycin resistance cassette *aac*(*3*)*IV*, flanked by *lox*P sites (apra-*lox*P), was subsequently cloned into the engineered *Xba*I sites between the flanks to create *rokL6* knock-out construct pKO-*rokL6*. In the same way, the flanking regions of sco1448 were cloned into pWHM3 with the apra-*lox*P to produce sco1448 knock-out vector, pKO-1448. The knock-out constructs were introduced into *S. coelicolor* M145 or its *nagB* mutant. For clean gene knock-out mutants, the apramycin resistance cassette was excised by introduction of pUWLcre, which expresses Cre recombinase (34, 35). The correct recombination event in each of the knock-out mutants was confirmed by PCR.

For *rokL6* complementation, the entire coding region of *rokL6* with its own promoter was amplified from the *S. coelicolor* chromosome. The PCR product was digested with *Xba*I/*Eco*RV and then inserted into pSET152 (36) to obtain *rokL6*-complemented vector pCOM-*rokL6*. This construct was introduced into *nagB*-*rokL6* double mutant Δ*nagB*Δ*rokL6*, and the successfully complemented strains Δ*nagB*Δ*rokL6-*C were selected by apramycin. The empty pSET152 was conjugated in Δ*nagB*Δ*rokL6* to produce Δ*nagB*Δ*rokL6-*E, which was used as a control strain.

For SCO1448 overexpression, a 1212-bp DNA fragment containing sco1448 was amplified and ligated into pSET152 with the 63-bp highly efficient promoter P30, namely the promoter SP30 with the 20-bp ribosomal binding site RBS15 (37), to produce SCO1448-overexpressing vector pOE-1448. The pOE-1448 construct was then introduced into Δ*nagB* to obtain the SCO1448 overexpression strain Δ*nagB*SCO1448-O.

### FLAG_3_ tag knock-in by CRISPR

To express FLAG-tagged RokL6 in *S. coelicolor*, the 3×FLAG sequence (FLAG_3_) was fused to the end of original copy of *rokL6* on genome using codon optimized CRISPR-Cas9 system (38) as described previously (39). Briefly, the spacer sequence (5’- GGCGCCGCGGGCTACCGCTC- 3’) specific to *rokL6*, located at the end of the coding region, was inserted into the pCRISPomyces-2 plasmid from *Bbs*I sites. Next, the 2263-bp template containing FLAG_3_ for homology-directed repair (HDR) was made and inserted into the spacer-containing plasmid, resulting in a *rokL6*- FLAG_3_ knock-in construct designated as pKI-FLAG_3_. Mutagenesis was done according to the previous study (39). After conjugation of pKI-FLAG_3_ to *S. coelicolor* A3(2) M145, ex-conjugants were patched on SFM agar plates with 20 ug/ml apramycin, then positive ex-conjugants were patched on antibiotic-free SFM agar plates and grown at 37°C.

Spores were collected and checked for loss of construct. Apramycin-sensitive strains were selected for spore collection, and their genomes were checked for desired recombination events. The successful *in situ* FLAG_3_ knock-in strain was identified as M145-*rokL6*-FLAG.

Additionally, *rokL6* with FLAG_3_ was ligated into pSET152 to generate pCOM-FLAG_3_ and introduced into the *nagB*-*rokL6* double mutant for evaluating RokL6-FLAG_3_ function *in vivo*.

### Heterologous expression and purification of His_6_-tagged RokL6 protein

For the heterologous expression of *S. coelicolor* RokL6 in *E. coli*, the 1197-bp *rokL6* coding region was amplified from *S. coelicolor* genomic DNA using primer pair RokL6-exp-F/RokL6-exp-R. The PCR fragment was ligated into pET-28a (+) from *Xho*I and *Nco*I sites, generating expression vector pEX-RokL6. The RoKL6 expression vector was transformed into *E. coli* BL21(DE3), and the expression of C-terminal His_6_-tagged RokL6 recombinant protein, RokL6- His_6_ was induced by the addition of Isopropyl β-D-1-thiogalactopyranoside (IPTG) at a final concentration of 1 mM when the cell density was reached around an optical density at 600 nm of 0.6, followed by overnight incubation at 16°C. Cells were harvested, washed, and disrupted in lysis buffer (40) by sonication. Soluble RokL6-His_6_ was purified from the supernatant using HisPur Cobalt Resin (Thermo fisher scientific; USA) and dialyzed against EMSA binding buffer.

### Chromatin immunoprecipitation sequencing (ChIP-Seq) of RokL6

ChIP-seq experiments were essentially carried out as described previously (39). In brief, *S. coelicolor* M145-*rokL6*-FLAG was grown on MM agar covered with cellophane disks, using mannitol with and without 50 mM GlcN as the carbon source. Mycelia were collected after 24 h (vegetative growth) and 48 h (sporulation). Mycelia were treated with phosphate-buffered saline (PBS) buffer containing 1% formaldehyde for 20 min to cross-link the DNA and protein. After thorough washing in PBS, mycelia were resuspended in lysis buffer (10 mM Tris-HCl pH 8.0, 50 mM NaCl, 15 mg/mL lysozyme, 1× protease inhibitor (Roche, Bavaria, Germany)) and incubated at 37°C for 20 min. After incubation, 0.5 mL IP buffer (100 mM Tris-HCl pH 8.0, 250 mM NaCl, 0.8% v/v Triton-X-100) was added to the mycelia samples, and chromosomal DNA was sheared to 100-500 bp fragments using the Bioruptor Pluswater bath sonication system (Diagenode, Liège, Belgium). Lysates were incubated with 40 μL Anti-FLAG M2 affinity gel (cat A2220, Sigma-Aldrich, St. Louis, US) and incubated at 4°C overnight. After centrifugation, the pellet and untreated total extracts (control) were incubated in 100 μL IP elution buffer (50 mM Tris-HCl pH 7.5, 10 mM EDTA, 1% m/v SDS) at 65°C overnight to reverse the cross-link. The beads were removed by centrifugation before DNA isolation by phenol-chloroform. The extracted DNA samples were then further purified with the DNA Clean & Concentrator kit (Zymo Research, California, US). The enriched DNA samples were sent to Novogene Europe (Cambridge, UK) for library construction and next-generation sequencing. ChIP-Seq data analysis was performed as described previously (39).

### RNA isolation, RNA sequencing (RNA-Seq) and quantity PCR (qPCR)

Spores of 10^7^ CFU of *S. coelicolor* M145 and the *rokL6* deletion strain Δ*rokL6* were grown on MM agar plates overlayed with cellophane discs, with either 1% (w/v) mannitol or 1% mannitol + 50 mM GlcN as the carbon sources. Biomass from two time points (24h on mannitol or 26h on mannitol with GlcN for vegetative growth phase, VEG; and 42h on mannitol or 44h on mannitol with GlcN for sporulation phase, SPO) was collected and snap-frozen in liquid N_2_. All samples were assessed as biological triplicates. After breaking the mycelia using a TissueLyser II (Qiagen, Venlo, The Netherlands), the RNA was extracted using a modified Kirby mix (31). The transcriptome sequencing library preparation and sequencing were outsourced to Novogen Europe (Cambridge, UK). Removal of rRNA from the samples was carried out using NEBNext Ultra directional RNA library prep kit (NEB, MA). Sequencing libraries were generated using the NEBNext Ultra RNA library prep kit for Illumina (NEB), and sequencing was performed on an Illumina NovaSeq 6000 platform. Raw data were cleaned using fastp v0.12.2 (41), and then mapped to the *S. coelicolor* M145 genome DNA (GeneBank accession AL645882.2) using bowtie2 v2.4.4 (42). Read counts for each gene were generated by featureCounts v2.0.1 (43). Values for transcripts per million (TPM) were generated using a custom python script. Differentially expressed genes and log2Foldchange were determined using DESeq2 v1.32.0 (44) with the data shrinkage function “apeglm” (45).

For quantitative PCR (qPCR), cDNA was synthesized using the iScript cDNA Synthesis Kit (Biorad, California, US). Briefly, qPCR was performed using the iTaq Universal SYBR green qPCR Kit (Biorad, California, US), and the program used was set as follows: 95°C 30 s; 40 cycles of 95°C 10 s, 60°C 30 s, plate reading; melting curve from 65°C to 95°C with 5 s per 0.5°C increment. The principal RNA polymerase σ factor encoding gene, *hrdB* (sco5820), was used as the internal control, and the qPCR data was analysed by CFX Manager software (version 3.1, Biorad, California, US), using ΔΔCq standard which is an implementation of the method described by (46).

### Electrophoretic Mobility Shift Assays (EMSAs)

EMSA were performed as described previously (39). Double stranded DNA probes (60-bp) were generated by gradually cooling reverse complemented single-strand oligonucleotides in 30 mM HEPES, pH 7.8, heating to 95°C for 5 min, then ramping to 4°C at a rate of 0.1°C/sec. The *in vitro* DNA-protein binding assays were performed in EMSA binding buffer (20 mM HEPES pH 7.6, 30 mM KCl, 10 mM (NH_4_)_2_SO_4_, 1 mM EDTA, 1 mM DTT, 0.2% Tween20). The binding reactions (10 μL), including 1 picomole DNA probe, 20 ng/μL BSA, and various concentration of RokL6-His_6_, were incubated at 30°C for 20 min. The reactions were then loaded on 5% non-denatured polyacrylamide gels and separated by electrophoresis. The gels were briefly stained with ethidium bromide and imaged using the Gel Doc imaging system (BioRad, California, US).

### Determination of transcriptional start sites (TSS)

Transcriptional start sites (TSS) of *rokL6* and sco1448 were determined by 5’ RACE using a 5’/3’ RACE Kit 2nd generation (Roche, California, US). Total RNA (2 μg) extracted from 48- hour cultures of *S. coelicolor* grown on solid minimal medium containing 1% mannitol was used for reverse transcription with gene-specific primers. The obtained cDNA was purified and an oligo(dA) tail added to the 3’ end by terminal transferase, followed by PCR amplification of the tailed cDNA with oligo(dT) anchor primer and gene-specific nested primers. Using the resulting PCR product (diluted 1000-fold) as a template, an additional round of PCR was performed with a more inner nested primer and an anchor primer provided in the kit to produce a single specific DNA band. The final PCR product was purified and sent for sequencing, and the TSS was determined as the first nucleotide following oligo(dA).

### Promoter activity test

The promoter regions of *rokL6*, sco1448, *nagKA* (12) and P30 (37) were ligated with a gene expressing enhanced green fluorescent protein (eGFP), and integrated into pSET152 via the *Xba*I/*Eco*RV sites, generating four recombinant vectors: pEGFP-*rokL6*, pEGFP-1448, pEGFP- *nagKA* and pEGFP-P30. The recombinant constructs were then introduced into *S. coelicolor* and *rokL6* deletion mutant Δ*rokL6* to generate the following strains: M145-P*rokL6*-eGFP, M145-P1448-eGFP, M145-P*nagKA*-eGFP, M145-P30-eGFP, and Δ*rokL6*-P*rokL6*-eGFP, Δ*rokL6*-P1448-eGFP, Δ*rokL6*-P*nagKA*-eGFP, Δ*rokL6*-P30-eGFP. The *in vivo* activities of all the promoters were evaluated by comparing the transcription level of the gene for eGFP via confocal microscopy and confirmed by qPCR.

### Confocal imaging

Sterile coverslips were inserted into MM with 1% mannitol agar plates at an angle of 45°, and spores of eGFP-harboring strains were inoculated at the intersection angle and incubated at 30°C for 48 h. The mycelium was scrapped off from coverslips into a drop of water, and then imaged with a TCS SP8 confocal microscope (Leica) using a 63× oil immersion objective (NA: 1.40) (47). Image processing and the analysis of fluorescent intensity were performed using ImageJ (version 1.54d).

### Bioinformatics analysis

DNA and protein database searches were performed using the BLAST server of the National Centre for Biotechnology Information (http://www.ncbi.nlm.nih.gov) and the *S. coelicolor* genome page services (http://strepdb.streptomyces.org.uk). Polypeptide sequences were aligned using Clustalw available at http://www.ebi.ac.uk/clustalw. Gene synteny analysis was performed using SynTax (48). To visualize the consensus sequence for the predicted binding site of RokL6, WebLogo (49) was used, and regulon predictions were done using PREDetector (50). The comparative analysis of the intergenic regions of *rokL6*/sco1448 and their orthologous pairs was performed by MEME (51).

### Data availability

Clean RNA-Seq reads and gene read-counts tables are available at GEO database (52) with accession number GSE234437 (https://www.ncbi.nlm.nih.gov/geo/query/acc.cgi?acc=GSE234437). Clean ChIP-Seq reads and binding region identification (peak calling) files are available at GEO database with accession number GSE234438 (https://www.ncbi.nlm.nih.gov/geo/query/acc.cgi?acc=GSE234438).

## RESULTS

### RokL6 is involved in GlcN metabolism

We previously showed that null mutants of *S. coelicolor nagB* are sensitive to high concentrations of GlcN, whereby spontaneous second-site mutations arise that allow the colonies to survive. Exploitation of this principle led to the identification of suppressor mutations in *nagA* and in a novel gene related to GlcN transport and metabolism (12, 24). Two independent suppressor mutants were obtained in the gene sco1447 when *S. coelicolor* Δ*nagB* was grown on GlcN, namely SMG1 and SMG38, and these mutations specifically alleviated the toxicity of exclusively on GlcN, but not GlcNAc (24, 53). Suppressor mutant SMG1 had sustained a single nucleotide insertion at nt position 26 within the coding region of sco1447, while SMG38 had a single nucleotide deletion at nt position 120 in sco1447. In both cases, the mutations resulted in a frame shift at the beginning of the gene, thereby preventing expression of the active protein. sco1447 encodes a ROK-family transcriptional regulator which we designated RokL6, based on the terminology that ROK-family proteins are named after the specific cosmid they are located on (25) within the ordered cosmid library that was used for the *S. coelicolor* genome sequencing project (54).

The observation that the mutations specifically alleviate toxicity of GlcN to *nagB* null mutants, but not that of GlcNAc, suggests that RokL6 plays a specific role in the control of GlcN metabolism. To test this, the *rokL6* single mutant Δ*rokL6* and *nagB-rokL6* double mutant Δ*nagB*Δ*rokL6* were generated by deleting *rokL6* in *S. coelicolor* M145 and in the previously published *nagB* null mutant (30), respectively. To create the mutants, the entire coding region of *rokL6* (nt positions +3 to + 1200) was replaced with the apramycin resistance cassette *aac*(*3*)*IV*, which was flanked by *lox*P sites. The *aac*(*3*)*IV* gene was subsequently excized from the genome using the Cre recombinase expressed from plasmid pUWLcre. The resulting *rokL6* single mutant and *nagB*-*rokL6* double mutant, Δ*nagB*Δ*rokL6*, was confirmed by PCR. In order to ascertain that the phenotypes were specifically caused by the deletion of *rokL6*, we genetically complemented the Δ*nagB*Δ*rokL6* mutant by expressing *rokL6.* For this, the -478/+1226 region of *rokL6*, harboring the entire gene and its promoter region, was amplified from the *S. coelicolor* chromosome and cloned into integrative vector pSET152 to obtain pCOM-*rokL6* (see Materials and Methods for details).

As expected, *S. coelicolor* M145 and its *rokL6* single mutant Δ*rokL6*, which has an intact copy of *nagB*, grew well on all media, while *nagB* mutants were unable to grow on MM with mannitol (1% w/v) and 5 mM GlcN or GlcNAc. The *rokL6*-*nagB* double mutant grew well on MM with 1% mannitol and 5 mM GlcN, but failed to grow when GlcNAc was used instead of GlcN (Fig. 1). Importantly, transformants expressing RokL6 in the Δ*nagB*Δ*rokL6* double mutant via the introduction of pCOM-*rokL6*, failed to grow, similar to the *nagB* mutant. These data strongly suggest that the deletion of *rokL6* was the sole reason why Δ*nagB*Δ*rokL6* double mutants could grow on GlcN (Fig. 1).

**Figure 1.**
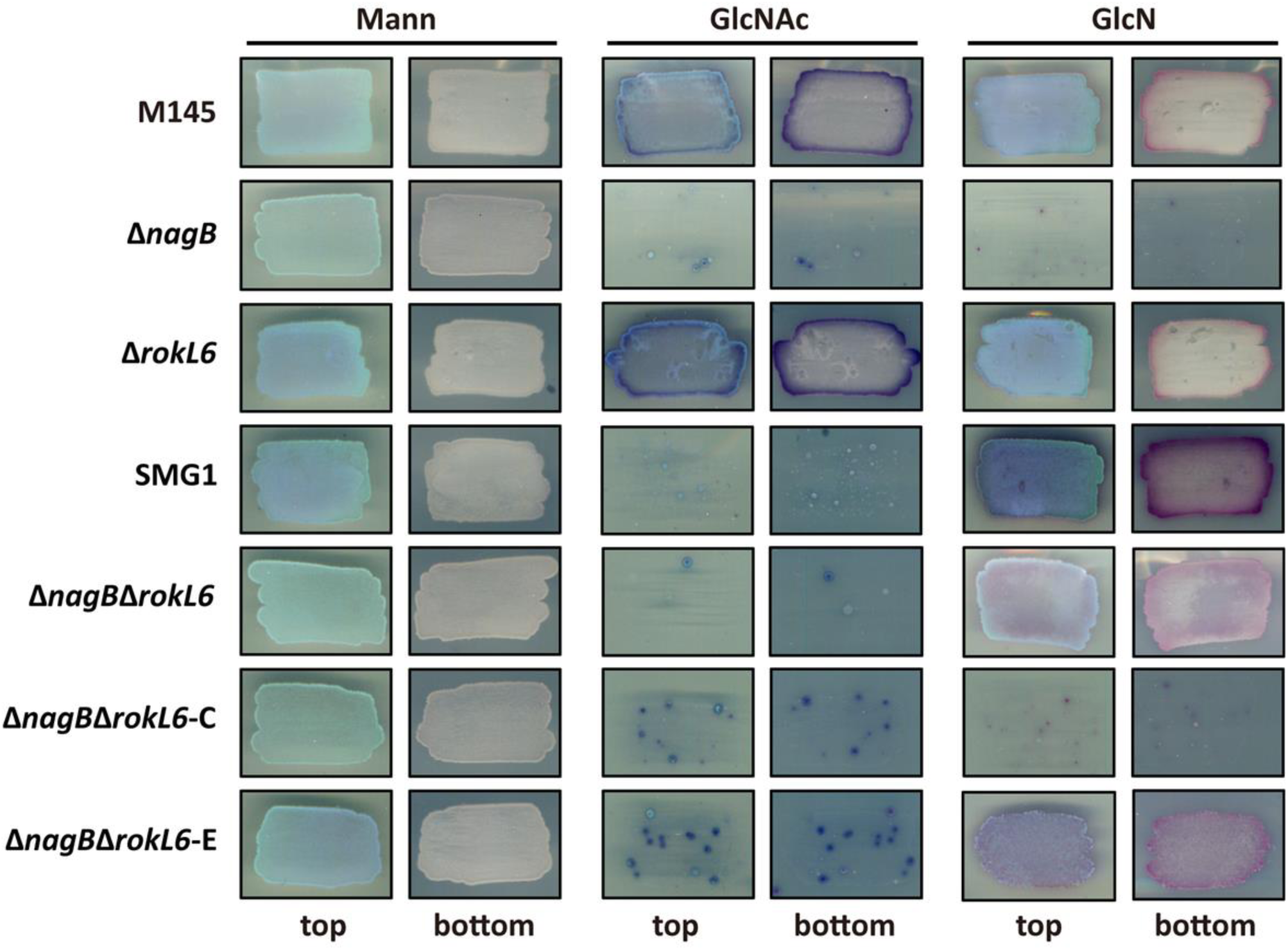
Sensitivity of *S. coelicolor* mutants to GlcN and GlcNAc. Spores (10^5^ CFU) of *S. coelicolor* M145 or its mutant derivatives were streaked onto minimal medium (MM) with either 1% (w/v) mannitol (Mann), 5 mM *N*-acetylglucosamine (GlcNAc) or 5 mM glucosamine (GlcN) and grown for 72 h at 30^°^C. Strains were *S. coelicolor* M145 (M145), its mutant derivatives Δ*nagB*, Δ*rokL6*, suppressor mutant SMG1, Δ*nagB*Δ*rokL6*, and the Δ*nagB*Δ*rokL6* mutant harboring either pCOM-*rokL6* (Δ*nagB*Δ*rokL6-*C) or empty vector pSET152 (Δ*nagB*Δ*rokL6-*E). Both top and bottom view are shown. Note that all strains without the gene *nagB* are sensitive to GlcNAc, while all strains lacking *rokL6* are resistant to GlcN.

### Conservation and gene synteny of *rokL6*

The gene *rokL6* (sco1447) from *S. coelicolor* consists of 1200 nucleotides (nt) and encodes a protein of 399 amino acids. RokL6 is characterized by an N-terminal winged helix-turn-helix (HTH) DNA-binding site which is found in various families of DNA-binding proteins and a sugar kinase domain annotated as a putative ROK-family regulator. Protein alignment shows a high amino acid sequence conservation of RokL6 in *Streptomyces* species (Fig. S1). Its genomic neighbors, sco1446 and sco1448, encode a putative integral membrane protein and a Major Facilitator Superfamily (MFS) transporter with unknown substrates, and the intergenic regions of *rokL6*-sco1446 and *rokL6*-sco1448 are 21 and 112 bp long, respectively (Fig. 2A). There is significant gene synteny for the genomic region surrounding *rokL6* and its orthologues in *Streptomyces*. In particular, sco1448 and its homologs, often lie divergently transcribed from the corresponding ROK regulators encoding genes (Fig. 2B). The *rokL6*-sco1448 unit is also identified in many *Kitasatospora* species (Fig. S2).

**Figure 2.**
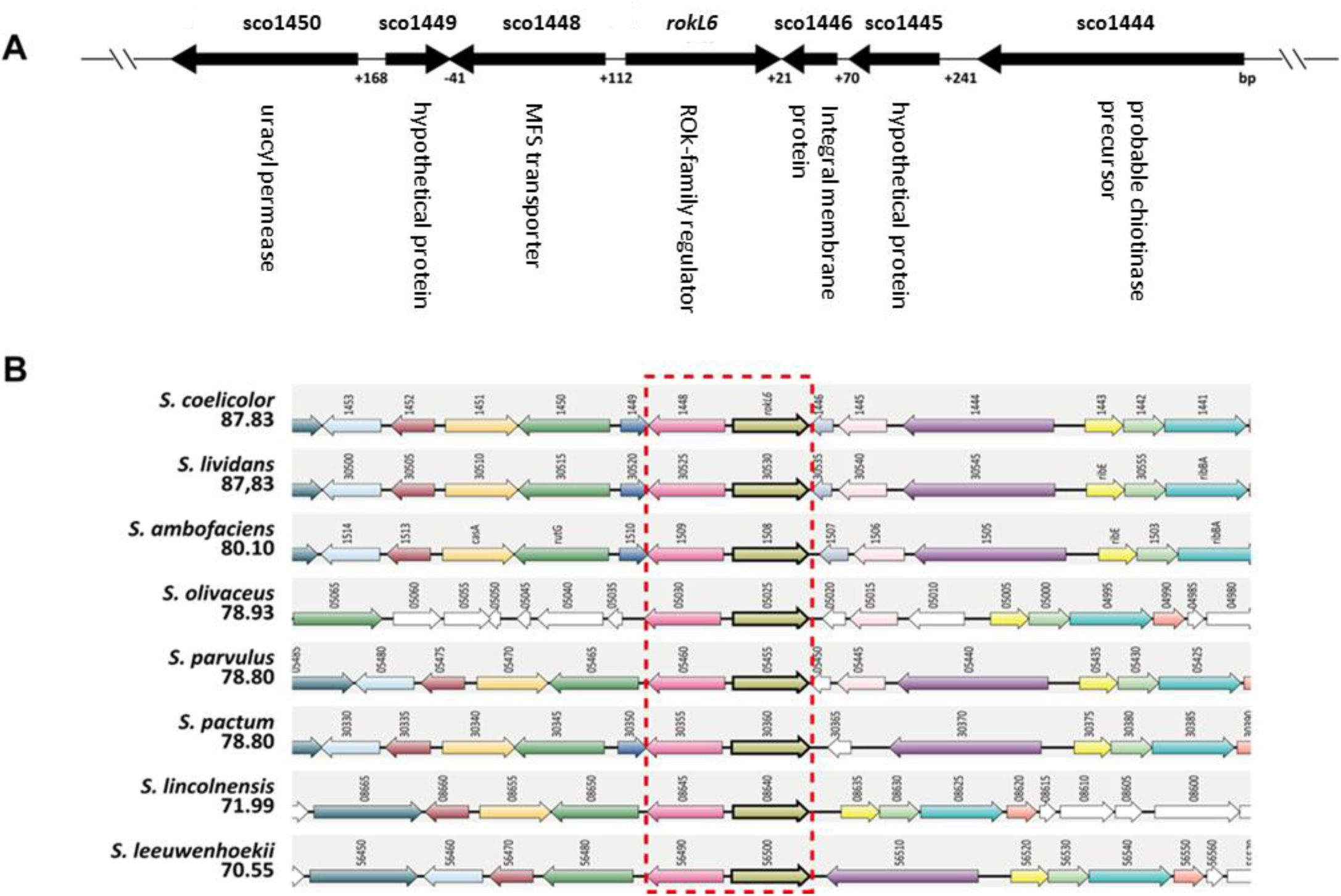
Genetic organization and gene synteny around *rokL6*. (A) Genetic organization of the genomic region around *rokL*6. All open reading frames (ORFs) are depicted by solid arrows with their predicted gene products shown underneath; (B) Gene synteny of the region around *rokL6* in selected *Streptomyces* species. Synteny analysis was performed by Synttax (scores are given). Homologous genes are presented in the same colors. Homologs of *rokL6* and its neighbor sco1448 are indicated in the dashed red box.

Given the importance of ROK-family regulators in the control of sugar utilization (26) and the likelihood of the involvement of RokL6 in GlcN metabolism, we investigated its potential role in GlcN utilization by measuring the dry weights at different time points to compare the growth patterns of *S. coelicolor* M145 and its *rokL6* mutant in NMMP containing 1% GlcN as the sole carbon source. The mutant grew well in NMMP supplemented with either 1% glucose or with 1% GlcN (Fig. S3).

### Transcriptome analysis of the *rokL6* mutant

To obtain further insights into the regulon of RokL6 and its relationship to GlcN metabolism, RNA-Seq analysis was performed using RNA extracted from *S. coelicolor* M145 and its *rokL6* mutant grown on MM agar with either mannitol or mannitol + 50 mM GlcN as the carbon sources. Biomass was harvested at two time points corresponding to vegetative growth or sporulation. In total 29 genes were significantly differentially expressed under at least one of the conditions analysed, using an adjusted *p*-value < 0.01 and a log2 fold change >2 or < -2 as threshold (Table S1). Genes sco1446 and sco1448-sco1450, which flank *rokL6,* were up- regulated in the *rokL6* mutant under all tested conditions (Fig. 3). In addition, sco0476 (for an unknown ABC transport protein) was down-regulated at 24h. In the absence of GlcN, transcription of the *pstSCA* operon (sco4140-4142), which is part of the PhoP regulon and related to the transport of inorganic phosphate (55), was significantly up-regulated (> 4- fold), while sco5338 (probable regulatory protein) and sco5339 (probable plasmid transfer protein) were down-regulated in Δ*rokL6* during sporulation (Fig. 3A). In the presence of GlcN, the putative ABC transporter genes sco3704-sco3706 were up-regulated in Δ*rokL6* during sporulation in the presence of GlcN; transcription of genes in the biosynthetic gene cluster for carotenoid biosynthesis (56), including *crtE* (sco0185), *crtI* (sco0186), *crtB* (sco0187), *crtY* (sco0191), *litQ* (sco0192) and the regulatory gene *litR* (sco0193), were all strongly down-regulated in the *rokL6* mutant during vegetative growth (Fig. 3B). Indeed, compared to the wild-type M145, the production of carotenoids was significantly reduced in Δ*rokL6* in the presence of GlcN when plates were incubated in the light (Fig. S4), suggesting that RokL6 indeed plays a role in the control of carotenoid biosynthesis in *S. coelicolor*.

**Figure 3.**
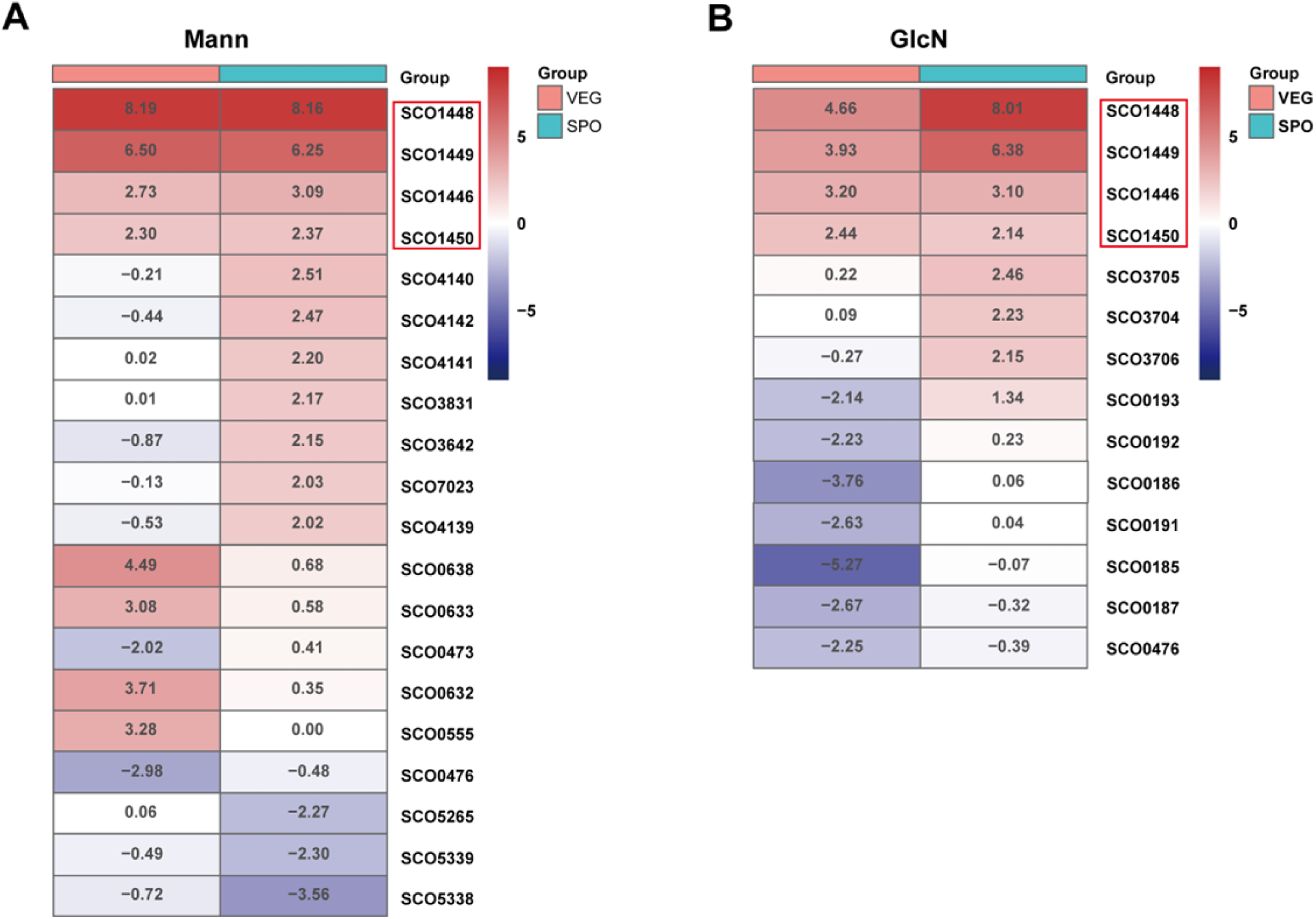
Heat maps of genes differentially expressed between the *rokL6* mutant and its parent *S. coelicolor* M145. Transcription patterns (expressed as log2 fold changes Δ*rokL6*/wild-type) are presented for genes differentially expressed when grown on MM with mannitol (A) or on MM with mannitol and GlcN (B). Only genes with an adjusted *p*-value less than 0.01 are shown. Navy, down-regulated (log2 fold change < -2) and brick red, up-regulated (log2 fold change > 2) in the *rokL6* mutant; intermediate log2 fold changes represented in white. Genes significantly differentially expressed under all growth conditions are highlighted with red boxes. VEG, vegetative growth phase; SPO, sporulation.

### RokL6 specially binds to the intergenic region of *rokL6* and sco1448

To identify direct targets of RokL6 *in vivo*, we performed Chromatin Immuno-Precipitation combined with sequencing (ChIP-Seq). For this, we constructed a strain expressing RokL6 with a FLAG_3_ tag, as described in the Materials and Methods section. We used CRISPR-Cas9 technology to construct strain *S. coelicolor* M145-*rokL6*-FLAG, which has an engineered in-frame 3×FLAG epitope fused *in situ* and before the stop codon of *rokL6*. The functionality of RokL6-FLAG_3_ was verified by introducing pCOM-FLAG into Δ*nagB*Δ*rokL6*. Indeed, while Δ*nagB*Δ*rokL6* mutants grow well on MM with mannitol and GlcN, Δ*nagB*Δ*rokL6*-FLAG expressing RokL6-FLAG_3_ failed to grow (Fig. S5), showing that the RokL6-FLAG_3_ was indeed expressed and active. ChIP-Seq analysis was performed with M145-*rokL6*-FLAG using cultures grown for 24 h and 48 h on MM with mannitol and MM with mannitol and GlcN. Under all conditions tested, only one major binding event was observed in all of the samples, namely to the intergenic region of *rokL6* and sco1448 (Fig. 4A). Binding of RokL6 to the intergenic region shared between *rokL6* and sco1448 was further verified *in vitro* by electromobility shift assays (EMSAs). For this, C-terminally 6×His-tagged RokL6, RokL6-His_6_, was expressed and purified from *E. coli* BL21(DE3). Compared to the probe for negative control, *hrdB* promoter (*hrdB*-P), which exhibited no retardation in the presence of RokL6, the result shows that RokL6 caused a strong retardation on the probe corresponding to the promoter region of *rokL6*-sco1448 (sco1448-P). Specifically, almost all sco1448-P was bound when 1 μM RokL6-His_6_ was added in the EMSA (Fig. 4B).

**Figure 4.**
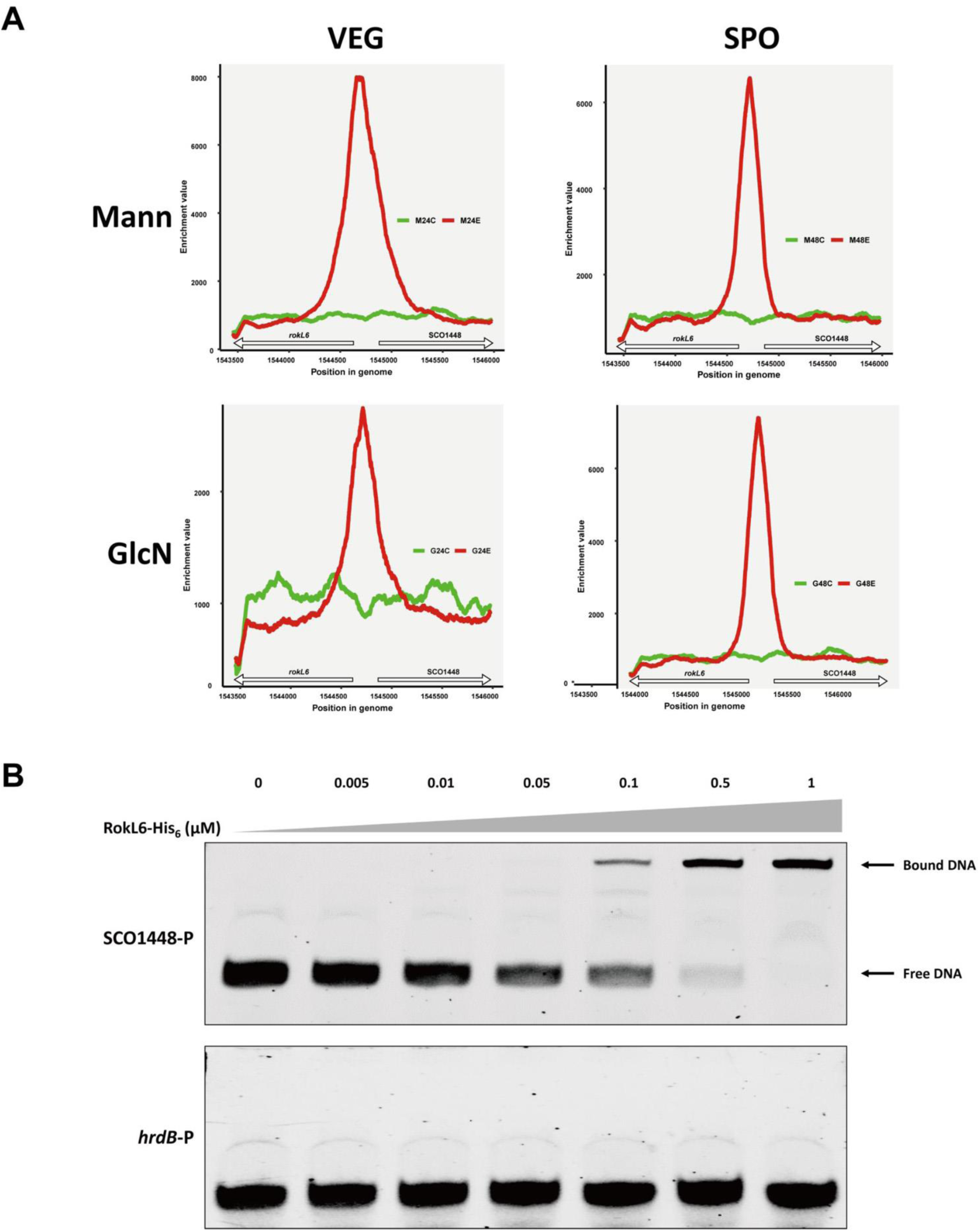
Identification of RokL6 binding sites *in vivo* and *in vitro*. (A) Enrichment peaks of RokL6 by ChIP-Seq. Red line, ChIP sample; green line, input chromosomal DNA used as a negative control. Genes flanking the peak summits are indicated. Numbers on the *X*-axis indicate genomic positions. Samples were collected on mannitol (Mann) and mannitol with GlcN (GlcN) at two time points, vegetative phase (VEG) and sporulation phase (SPO). (B) EMSAs to establish direct binding in vitro. The binding of RokL6 to the intergenic sequence between *rokL6* and sco1448 (sco1448-P) was tested by EMSAs, with the *hrdB* promoter (*hrdB*-P) sequence as the negative control. Concentrations given in μM.

### RokL6 binds to the inverted repeat *rokL6*-IR

Due to the high conservation in terms of both amino acid sequence and gene synteny of *rokL6* and sco1448 with their orthologous, we reasoned that the orthologs of RokL6 may be autoregulators. To identify a putative RokL6 binding consensus, the intergenic DNA sequence of *S. coelicolor rokL6* and sco1448, and 10 additional orthologous gene pairs from other streptomycetes (Table 3) were analysed for the presence of conserved motifs using MEME.

**Table 3.**
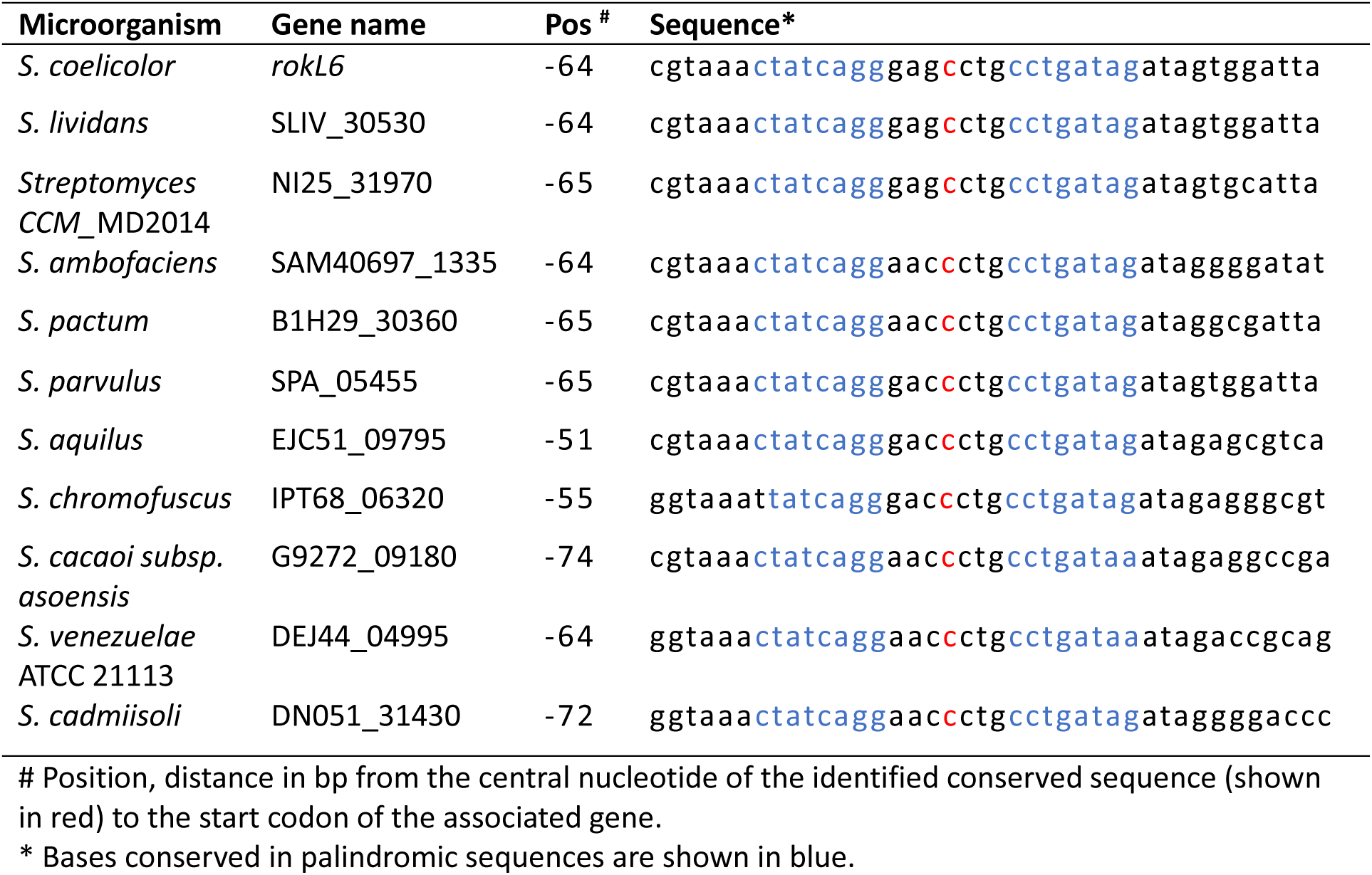
Alignment of palindromic sequences found upstream of genes encoding RokL6 and its orthologs in streptomycetes.

This identified a highly conserved inverted repeat sequence of 23 nucleotides in all intergenic regions (Fig. 5A). The consensus of the motif was C(T)TATCAGG - 7 nt - CCTGATAG(A), which contains an inverted repeat designated as *rokL6*-IR, a primary candidate for the RokL6 binding site.

**Figure 5.**
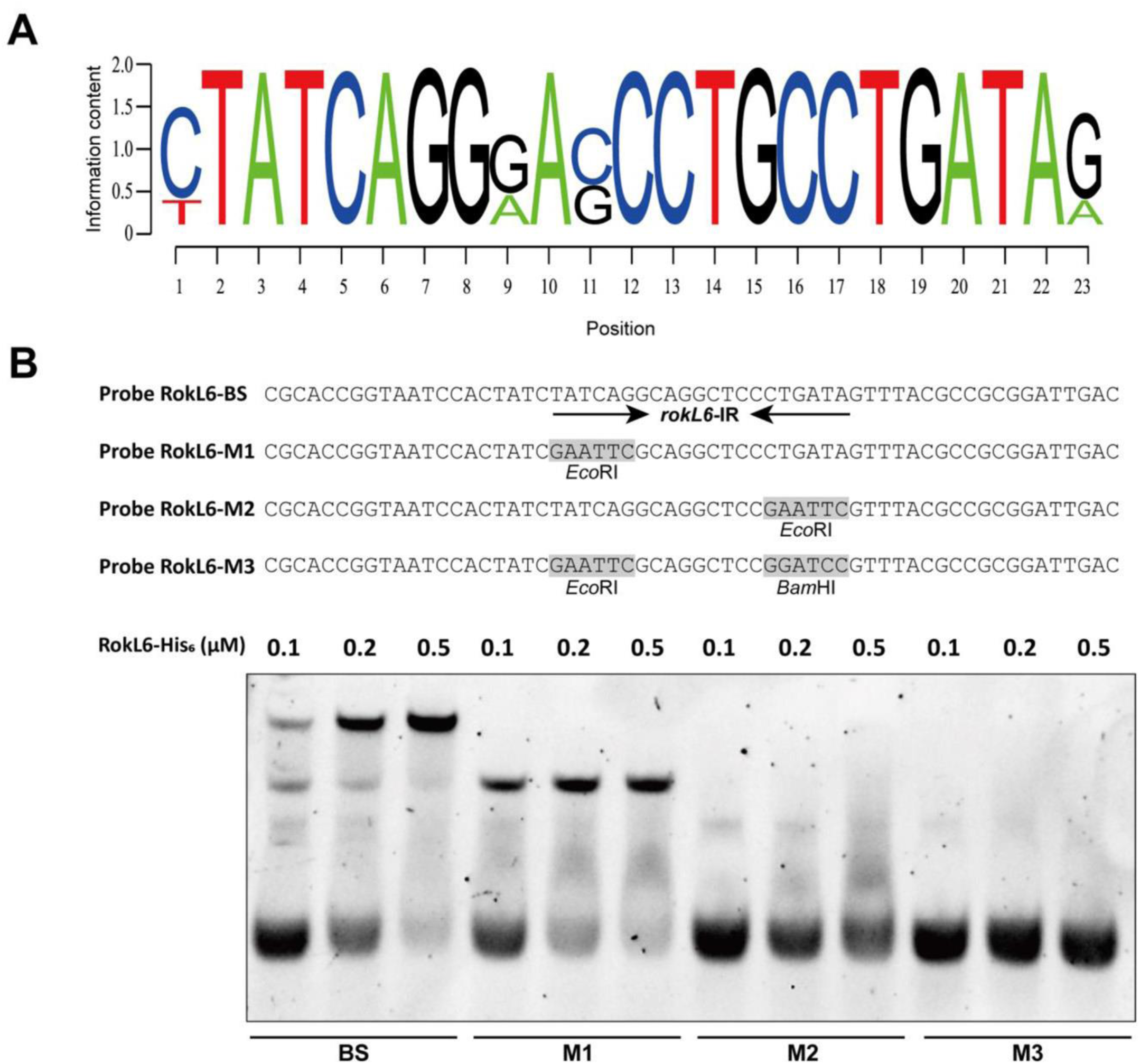
Identification of the RokL6 binding motif. (A) Sequence Logo representation of consensus sequence of RokL6 binding sites (analyzed by WebLogo). (B) RokL6 binding sites confirmation by EMSA. EMSAs of RokL6-His_6_ with four probe sequences (RokL6-BS, RokL6- M1, RokL6-M2, RokL6-M3) were tested. The palindromic sequences (*rokL6*-IR) in Probe RokL6-BS are indicated by arrows. For each probe tested, the concentrations of RokL6-His_6_ were 0.1, 0.2, and 0.5 μM.

To verify if this is indeed a *bona fide* RokL6 binding site, EMSAs were performed using 60 bp probes. These probes contained either the intact *rokL6*-IR sequence (probe RokL6-BS) or mutant versions in which one or two regions of *rokL6*-IR were replaced by *Eco*RI or *Bam*HI restriction sites (probe RokL6-M1 to M3). Compared to RokL6-BS, which was bound at all tested concentrations (0.1, 0.2, and 0.5 μM) of RokL6-His_6_, the binding signals of RokL6-His_6_ to the mutated probes RokL6-M1 and RokL6-M2 was weakened, and to RokL6-M3 was fully abolished (Fig. 5B). Thes data show that the inverted repeat sequence *rokL6*-IR is indeed an essential part of the RokL6 binding site.

To determine if RokL6 has other binding sites on the *S. coelicolor* genome, the sequence of *rokL6*-IR was used to scan the *S. coelicolor* genome using PREDetector algorithm (50). In addition to the promoter regions of *rokL6* and sco1448, one putative motif was identified with relative high score upstream of sco0137 encoding a PTS EII with unknown substrate (Table S2). EMSAs showed that binding between RokL6 and the promoter of sco0137 (sco0137-P) *in vitro* was very weak (Fig. S6A). The putative PTS transporter EII genes sco0137 and sco0136 are co-expressed from a single transcriptional unit; to see if the operon would be the major GlcN transporter, a mutant was created in both the wild-type strain *S. coelicolor* M145 and in the *nagB* mutant. For this, the region from nt position +10 of sco0137 to nt position + 1542 of sco0136 was replaced by *aac*(*3*)*IV* (see Materials and Methods). Deletion of sco0136-sco0137 in the *nagB* mutant did not relieve GlcN toxicity (Fig. S6B). This strongly suggests that sco0136-sco0137 are at least not solely responsible for GlcN transport in *S. coelicolor*.

### Identification of the transcription start sites (TSS) for *rokL6* and sco1448 and repression by RokL6

5’ rapid amplification of cDNA ends (5’ RACE) was applied to identify the promoters of both *rokL6* and sco1448. Fragments of 5’ cDNA of *rokL6* and sco1448 were PCR-amplified from the *S. coelicolor* genome and sequenced (Fig. S7). The TSS of *rokL6* was localized to a G located 67 nt upstream of the *rokL6* translation start codon (TSC), and the sco1448 TSS was mapped to a G located 27 nt upstream of the sco1448 TSC (Fig. 6A). The RokL6 binding site *rokL6*-IR encompasses nt positions -7 to +14 relative to the *rokL6* TSS, which is immediately downstream of the putative -10 region of the *rokL6* promoter. Suggestively, the *rokL6*-IR is located exactly between the -10 and -35 sequences of the sco1448 promoter. This places the RokL6 binding site in the ideal position to repress both genes at the same time. Binding of the repressor close to the -10 or -35 consensus boxes for the RNA polymerase sigma factor is common, as it interferes with the recognition, binding of RNA polymerase, and the formation of the transcriptional initiation complex (57). This suggests that RokL6 represses the transcription of both sco1448 and *rokL6* by binding to the core promoter regions to prevent their transcription from proceeding properly.

**Figure 6.**
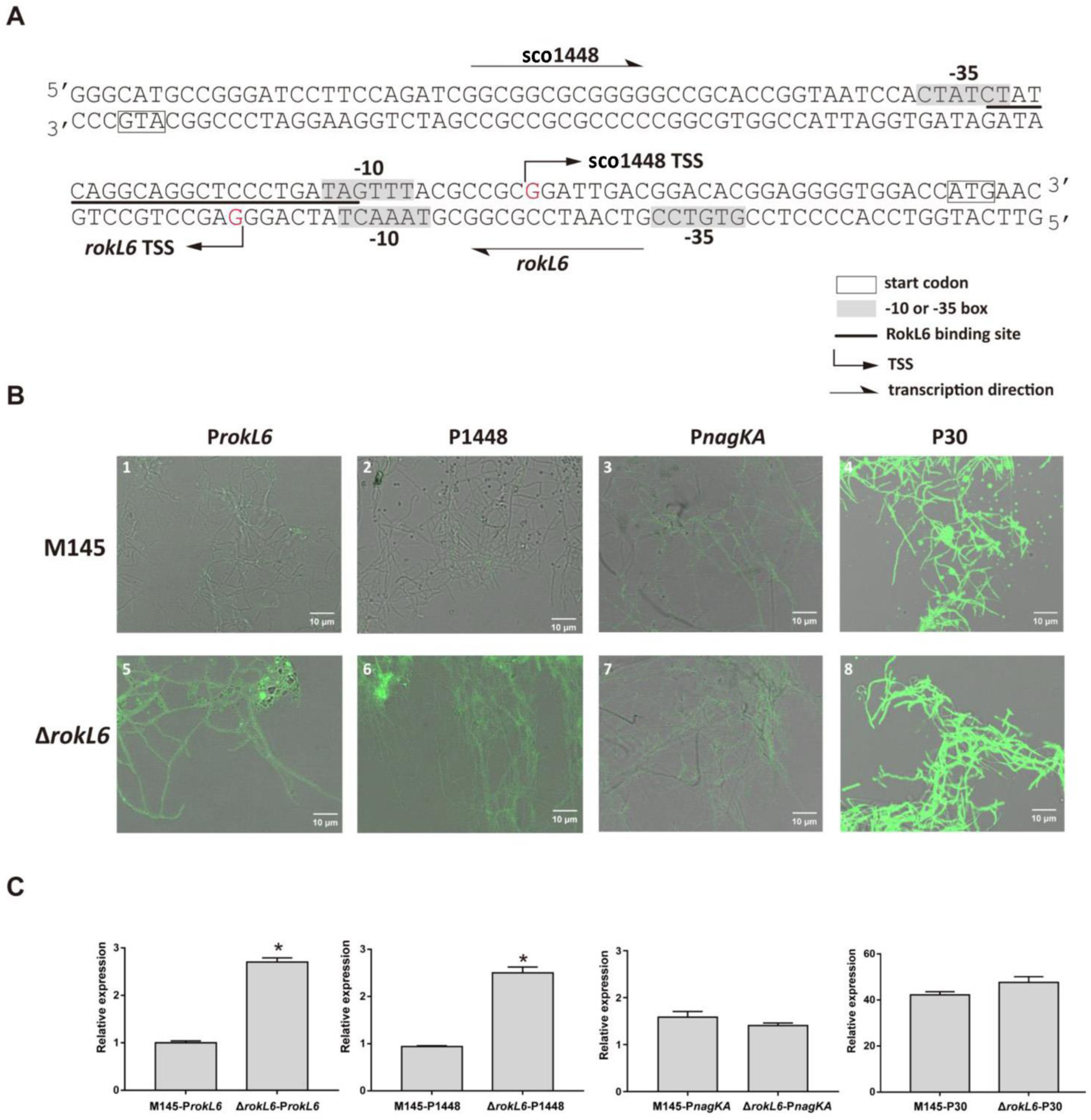
RokL6 acts as transcriptional repressor of sco1448 and as auto-repressor. (A) Nucleotide sequences of *rokL6* and sco1448 promoter region and RokL6-binding site. Number, distance (nt) from the respective translation start codon (TSC); bent arrows, transcription start sites (TSS); boxes, start codons; grey shading, predicted -10 or -35 boxes based on 5’RACE (Fig. S7); underlined, RokL6 binding site; straight arrow, direction of transcription. (B) Fluorescence intensity of eGFP measured based on confocal fluorescence micrographs. Mycelia of the strains transformed by eGFP with different promoters were imaged using confocal microscopy. Strains analysed as indicated: 1, M145-P*rokL6;* 2, M145- P1448; 3, M145-P*nagKA*; 4, M145-P30; 5, Δ*rokL6-*P*rokL6*; 6, Δ*rokL6*-P1448; 7, Δ*rokL6*-P*nagKA*; 8, Δ*rokL6*-P30. (C) Relative transcription levels of eGFP. Transcription levels of for eGFP gene expressed from different promoters (P*rokL6*, P1448, P*nagKA* and P30) in M145 and Δ*rokL6* were measured by qPCR analysis. Relative transcription levels were normalized to the transcription level of the gene for eGFP with P*rokL6* in M145, which was set as 1. Data were calculated from triplicate biological experiments and presented as mean ± SD (* *P* <0.05).

To determine if this is indeed the case, promoter activity assays were performed using eGFP as the reporter gene cloned in integrative vector pSET152, using promoters P*rokL6* (*rokL6* promoter) and P1448 (sco1448 promoter), with the DasR-controlled *nagKA* promoter (P*nagKA)* and P30 as the controls. The resulting vectors pEGFP-*rokL6*, pEGFP-1448, pEGFP-*nagKA* and pEGFP-P30 were introduced into *S. coelicolor* M145 and Δ*rokL6*. The recombinant strains were grown on MM agar with 1% mannitol and analysed for eGFP expression levels by florescence microscopy and qPCR. When eGFP was expressed from either the *nagKA* promoter or the artificial promoter P30, fluorescence was unchanged between M145 and Δ*rokL6*. Conversely, when expressed from either P*rokL6* or P1448, eGFP levels were significantly higher in Δ*rokL6* as compared to the parental strain M145 (Fig. 6B). This result was confirmed by qPCR for the transcription of the gene for eGFP, with enhanced expression of eGFP in the *rokL6* mutant as compared to the parent, while again no significant differences in expression levels were observed for either P*nagKA* or P30.

However, the transcript levels of the gene for eGFP from both P*rokL6* and P1448 were significantly up-regulated in Δ*rokL6* (Fig. 6C), suggesting that RokL6 indeed represses the transcription of both sco1448 and at the same time acts as an autoregulator.

### Overexpression of transporter SCO1448 relieves GlcN toxicity in *nagB* mutants

The data above demonstrate that putative transporter protein SCO1448 is the primary target of RokL6. We therefore hypothesized that SCO1448 may function as an exporter of a toxic metabolic intermediate that accumulates in *nagB* mutants grown on media containing GlcN. Inactivation of the repressor gene *rokL6* results in enhanced expression of SCO1448, alleviating GlcN toxicity, presumably by exporting a toxic intermediate. If this is indeed the case, expressing sco1448 from a RokL6-independent (constitutive) promoter should have the same effect as deleting *rokL6*. To test this, sco1448 was placed under the control of the strong and constitutive P30 promoter (see Materials and Methods for details), and introduced into Δ*nagB* to obtain SCO1448 over-expression strain, Δ*nagB*SCO1448-O. As a control, we used empty plasmid pSET152, to obtain Δ*nagB*-E. Transcription levels of sco1448 were measured, showing that in both the de-repressed Δ*nagB*Δ*rokL6* and the overexpressing Δ*nagB*SCO1448-O strains, SCO1448 was highly expressed (Fig. 7A).

**Figure 7.**
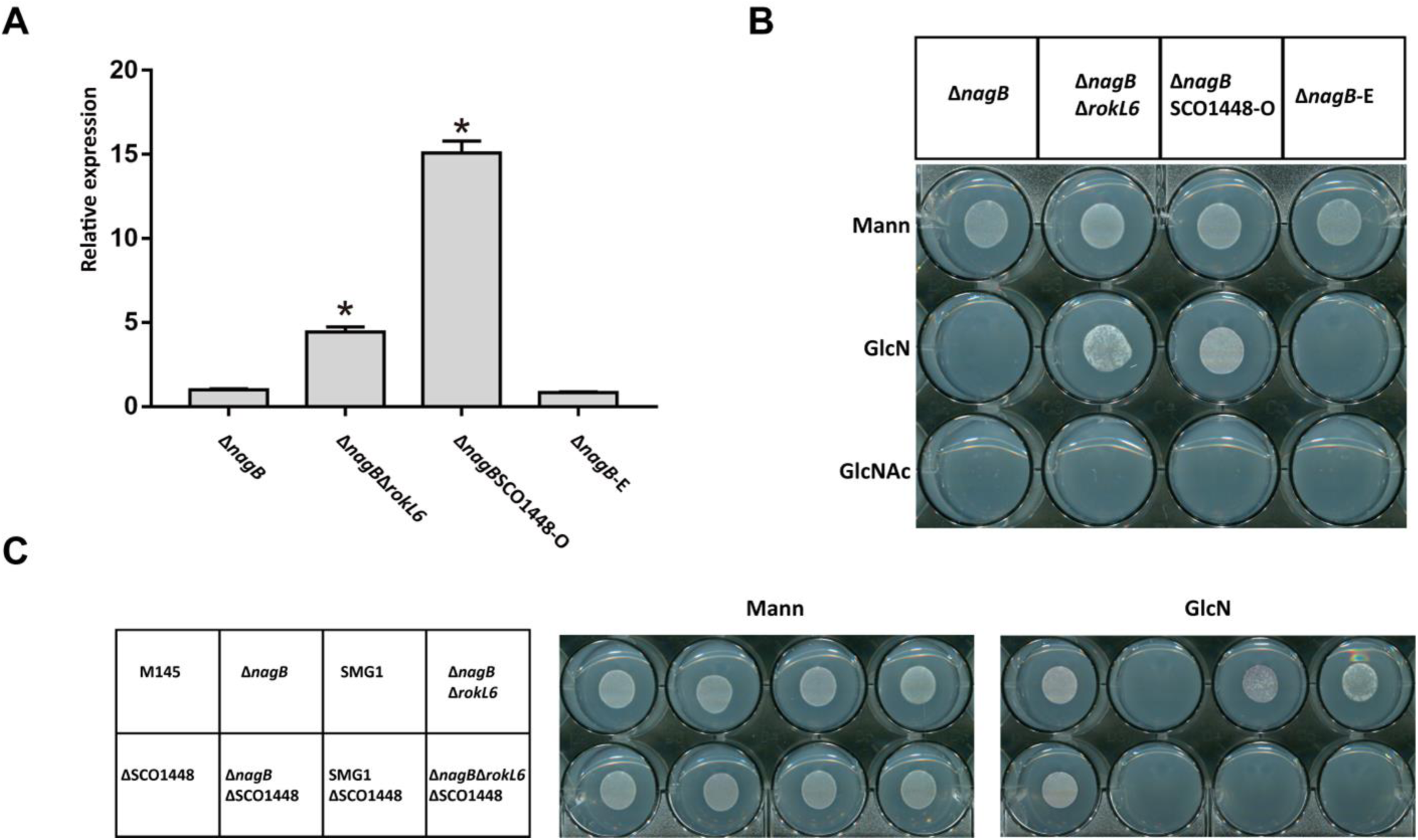
Constitutive expression of sco1448 relieves toxicity of *nagB* mutants to GlcN. (A) sco1448 transcription levels analyzed by qPCR of the strains as follows, Δ*nagB*, Δ*nagB*Δ*rokL6*, Δ*nagB* with overexpressed sco1448 (Δ*nagB*SCO1448-O), and Δ*nagB* complemented with empty pSET152 (Δ*nagB*-E). qPCR data were calculated from three independent experiments and presented as mean ± SD (* *P* < 0.05). (B) Growth of Δ*nagB*SCO1448-O on MM with GlcN(Ac). Spores of Δ*nagB*, Δ*nagBΔrokL6*, Δ*nagB*SCO1448-O and Δ*nagB*-E, as indicated on the top, were spotted on MM with mannitol (Mann), MM with 1% mannitol and 5 mM GlcN (GlcN), and MM with 1% mannitol and 5 mM GlcNAc (GlcNAc) to determine their growth. (C) Growth of sco1448 knock-out strains on GlcN. For media see (B).

Importantly, RokL6-independent expression of SCO1448 fully alleviated toxicity of *nagB* mutants to GlcN, while Δ*nagB* or Δ*nagB* with the empty plasmid were still sensitive to GlcN (Fig. 7B).

Finally, we evaluated the viability of sco1448 knock-out mutants in the presence of GlcN. For this, the coding regions of sco1448, spanning nucleotides +6 to +1023 (relative to the transcriptional start site), were substituted with *aac*(*3*)*IV*, in *S. coelicolor* M145, Δ*nagB*, SMG1 (Δ*nagB* with a suppressor mutation in *rokL6*), and Δ*nagB*Δ*rokL6*. Homologous recombination of the gene was achieved by introduction of vector pKO-1448, which was constructed by cloning the resistance cassette between the upstream and downstream flanking regions of sco1448 in the unstable multi-copy plasmid pWHM3. Correct recombination events were confirmed by resistance to apramycin and sensitivity to thiostrepton (for loss of the plasmid) and by PCR, thus obtaining sco1448 knock-out mutants, Δsco1448, Δ*nagB*Δsco1448, SMG1Δsco1448, and Δ*nagB*Δ*rokL6*Δsco1448. As expected, when sco1448 was deleted in suppressor mutant SMG1 or in Δ*nagB*Δ*rokL6*, neither strain could grow in the presence of GlcN (Fig. 7C). This again provides compelling evidence that sco1448 is solely responsible for alleviating the toxicity of GlcN to *nagB* mutants. Taken together, our data show that RokL6 directly represses the expression of sco1448, and that in turn, sco1448 is responsible for alleviating GlcN toxicity to *nagB* mutants. It is likely that SCO1448 exports toxic metabolic intermediates derived from GlcN- 6P that accumulates in *nagB* mutants of *S. coelicolor* when grown on GlcN.

## DISCUSSION

The aminosugar *N*-acetylglucosamine (GlcNAc) is a preferred nutrient for *Streptomyces* and also acts as a signaling molecule for the nutritional status of the environment. While GlcNAc metabolism and transport in streptomycetes have been well studied, little is known about how GlcN is metabolized in *Streptomyces*. *S. coelicolor nagB* mutants, which cannot convert GlcN-6P into the glycolytic intermediate Fru-6P, fail to grow on either GlcNAc or GlcN, indicating that toxic intermediates are produced when GlcN-6P is not actively metabolized by NagB. We used this feature to select for suppressor mutants, whereby it is important to note that some *nagB* suppressors selected on GlcNAc fail to confer resistance to GlcN and *vice versa*. In this work we focused on the GlcN-specific gene *rokL6* (sco1447), mutation of which relieves GlcN toxicity in *nagB* mutants.

RokL6 is a ROK-family regulator, and members of this family of regulators often play a role in the control of sugar metabolism. Systems-wide analysis using RNA-Seq, qPCR, ChIP-Seq and EMSAs showed that RokL6 (SCO1447) directly represses the transcription ofsco1448, which is annotated as an MFS sugar transporter. SCO1448 plays a key role in alleviating GlcN toxicity in the absence of NagB activity. Up-regulation of sco1448 is sufficient to relieve GlcN toxicity to *S. coelicolor nagB* mutants, while conversely, deletion of sco1448 in *S. coelicolor* Δ*nagB*Δ*rokL6* or the suppressor mutant SMG1 abolishes the acquired GlcN resistance (Fig. 7). Thus, the key to GlcN resistance lies in the expression of SCO1448. Still, a lot is unclear about the *rokL6*-sco1448 gene pair. In particular, why would streptomycetes have a cryptic exporter to protect themselves from toxic intermediates related specifically to GlcN? The system is likely important, as its gene synteny and also (the location of) the RokL6 binding site are highly conserved in *Streptomyces* and its sister genus *Kitasatospora* (Fig. 2B and Table 3). The high conservation of the binding site predicts that expression of sco1448 is repressed by RokL6 in many if not all streptomycetes. While the biological significance it is not yet understood, the proteins likely play an important role in preventing the accumulation of excess toxic intermediates under specific growth conditions in the natural environment. How the repression by RokL6 is relieved in the cell and what the possible ligands are that control its DNA binding, which likely facilitates the derepression of sco1448, remains to be elucidated.

An important question is also what the exact nature is of the toxic molecule(s) that accumulate in *nagB* mutants, which are likely derived from GlcN-6P. Mutants that lack a functional NagA enzyme accumulate high levels of GlcNAc-6P, which is lethal in *E. coli* and *B. subtilis* (22, 23, 58). However, *S. coelicolor nagA* null mutants are able to grow in the presence of either GlcNAc or GlcN, and deletion of *nagA* from *nagB* null mutants of *S. coelicolor* also alleviate the toxicity of either aminosugar (24). However, there are no metabolic routes that point at a key role for NagA in GlcN metabolism. Why then would mutation of *nagA* prevent toxicity of not only GlcNAc but also GlcN in *nagB* mutants? This again shows that there are still significant gaps in our understanding of aminosugar metabolism in *Streptomyces*. This is in fact quite surprising for such an important central metabolic pathway.

ChIP-Seq and EMSA assays demonstrated that RokL6 binds directly to the palindromic *rokL6*-IR identified from the intergenic region of *rokL6* and sco1448, with the sequence 5’ -C(T)TATCAGG - 7 nt - CCTGATAG(A)- 3’. Furthermore, the precise transcription start sites for *rokL6* and sco1448 were identified by 5’ RACE, showing that the two genes are transcribed from overlapping promoters. RokL6 strategically binds to a site that is located precisely in between the -35 and -10 boxes for sco1448 and downstream of the -10 box for *rokL6*, allowing RokL6 to inhibit the transcription of both genes at the same time. The transcriptional repression of sco1448 and autoregulation of *rokL6* by RokL6 was further demonstrated by promoter activity assays and qPCR (Fig. 6), showing that RokL6 indeed inhibits the transcription of both genes.

The RokL6 binding site is distinct from operators characterized for ROK-family regulators in *e.g. E. coli* and Firmicutes. The operator consensus sequences of NagC and Mlc in *E. coli* or XylR in firmicutes are typically composed of two A/T-rich inverted repeats separated by a spacer of 5-9 bp (19, 59, 60). The binding consensus sequences of RokB from *S. coelicolor* and RokA from *Streptococcus pneumoniae* are also enriched with T and A at the 5’-end and 3’-end (61, 62), similar to CysR from *Corynebacterium glutamicum* (63). The binding target of CsnR from *S. lividans* shares some similarity with that of RokL6, containing the sequence 5’ -AGG - 7 nt - CCT- 3’ in the binding consensus. In contrast, the binding consensus of Rok7B7 of *S. avermitilis* is 5′ -<u>TT</u>K<u>A</u>MKHS<u>TT</u>S<u>A</u>V- 3′ and unrelated to that of RokL6 (19, 26).

ChIP-seq experiments identified a single binding site in all samples. Using that binding consensus as input, scanning the *S. coelicolor* genome using PREDetector revealed one additional sequence with similarity to the RokL6 consensus, namely upstream of the sco0137-sco0136 operon that encodes PTS transporter EIIC enzymes. Its control by RokL6 and the fact that in *E. coli* and *B. subtilis*, GlcN is transported via the PTS (64–66), suggested a possible role for the sco0137-sco0136 operon in GlcN transport. We tested whether inactivation of the operon would affect GlcN toxicity in *nagB* mutants, but this as not the case (Fig. S6B). Therefore, PTS transporter sco0136-sco0137 is at least not the only transporter for GlcN uptake in *S. coelicolor*, and more studies are required to understand how GlcN is internalized in streptomycetes.

In terms of indirect effects of the deletion of *rokL6,* besides sco1448 also the adjacent sco1446 and sco1448-sco1450 were upregulated in *rokL6* mutants. Other genes whose transcription was affected were those for transporters sco0476, sco3704-sco3706 and sco4140-sco4142, as well as for genes of the carotenoid biosynthetic gene cluster (BGC). Upregulation of the latter BGC was validated by enhanced pigmentation of mycelia grown in the light (Fig. S4). However, our experiments strongly suggest that the intergenic region between *rokL6* and sco1448 is the primary binding site for RokL6, and the other genes whose transcription was affected in *rokL6* mutants are likely controlled indirectly.

Our current understanding of aminosugar metabolism and transport in *Streptomyces* is shown schematically in Fig. 8, including possible functions for RokL6 and SCO1448. GlcN- 6P is derived either from the internalization of GlcN or the deacetylation of GlcNAc-6P. High accumulation of GlcN-6P and/or their metabolic derivatives is lethal to *S. coelicolor*, in line with observations in *E. coli* and *B. subtilis*. The MFS transporter SCO1448 likely serves to facilitate the export of toxic intermediate(s), and in turn, the transcription of sco1448 is repressed by RokL6 (Fig. 8). Constitutive expression of sco1448 does not relieve the toxicity of GlcNAc, suggesting that GlcNAc and GlcN toxicity are not caused by the same molecule(s).

**Figure 8.**
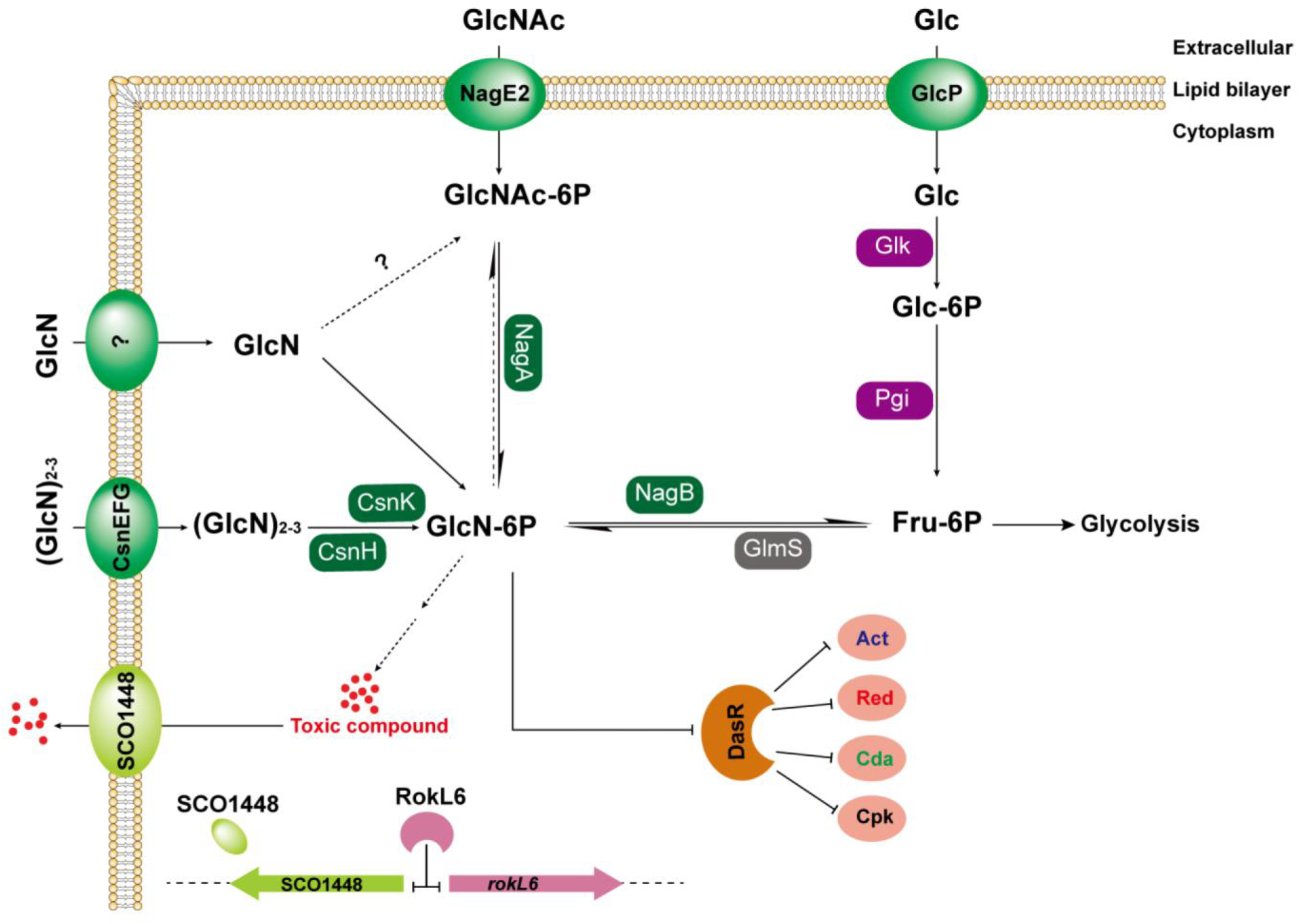
Model for aminosugar metabolism and the roles of RokL6 and SCO1448 in *S. coelicolor*. Metabolism of GlcNAc is well understood in *S. coelicolor*. GlcN-6P is an allosteric inhibitor of the global repressor DasR, which represses the biosynthesis of antibiotics. Based on our data, we propose that toxic substances are formed from GlcN metabolism, and particularly from GlcN-6P, which accumulates in GlcN-grown *nagB* mutants; these toxic substances are likely exported by the MFS transporter SCO1448. We show that this transporter is transcriptionally repressed by the ROK-family regulator RokL6, and that enhanced expression of sco1448 relieves GlcN toxicity. Metabolic routes are presented by black arrows with the enzymes as indicated/unknown routes by dotted arrows. Unknown transporters and enzymes are indicated by question marks. For details, see the main text. Abbreviations: GlcN, glucosamine; GlcNAc, *N*-acetylglucosamine; Glc, glucose; 6P, 6- phosphate; GlcP, glucose permease, Glk, glucokinase; Pgi, glucose-6-phosphate isomerase; GlmS, glucosamine-fructose-6-phosphate aminotransferase; Act, actinorhodin; Red, prodiginines; Cda, calcium-dependent antibiotic; Cpk, cryptic polyketide.

Taken together, this study characterizes the function of the ROK-family regulator RokL6, including its regulons, binding sites and its roles in the relief of GlcN toxicity in *S. coelicolor*. These findings shed new light on aminosugar sensitivity and on the control of aminosugar metabolism in *Streptomyces*, which plays a central role in their life cycle.

## ACKNOWLEDGMENTS

We thank E. de Waal for technical assistance. This work was supported by a fellowship from the Chinese Scholarship Council (CSC) to C. Li and by Advanced grant Community (grant number 101055020) by the European Research Council to GPvW.

