## Supplemental Figures and Tables for "Systems-wide analysis of the ROK-family regulatory gene *rokL6* and its role in the control of glucosamine toxicity in *Streptomyces coelicolor*"

#### Supplementary Tables

**Table S1.** Genes differentially expressed between the *rokL6* null mutant and its parent *S. coelicolor* M145 (wt).

| **Gene** | **Annotation** | **log2 fold change (Δ*rokL6*/wt)*** | | | |
|---|---|---|---|---|---|
| | | VEG-Mann | SPO-Mann | VEG-GlcN | SPO-GlcN |
| sco0555 | Membrane-bound oxidoreductase | 3.280 | 0.004 | -0.051 | 0.210 |
| sco0632 | RNA polymerase sigma factor SigK | 3.709 | 0.349 | -0.917 | 0.214 |
| sco0633 | Hypothetical protein | 3.079 | 0.578 | -0.668 | 0.178 |
| sco0638 | Lipoprotein | 4.490 | 0.683 | -0.184 | 0.8109 |
| sco1446 | Hypothetical protein | 2.73 | 3.093 | 3.199 | 3.099 |
| sco1448 | Transporter | 8.186 | 8.158 | 4.656 | 8.008 |
| sco1449 | Hypothetical protein | 6.500 | 6.247 | 3.926 | 6.382 |
| sco1450 | Uracyl permease | 2.300 | 2.372 | 2.444 | 2.135 |
| sco0473 | Solute-binding lipoprotein | -2.0157 | 0.4071 | -0.733 | -0.130 |
| sco0476 | ABC transporter ATP-binding protein | -2.981 | -0.475 | -2.250 | -0.392 |
| sco3642 | Hypothetical protein | -0.875 | 2.147 | -0.125 | -0.361 |
| sco3831 | Dehydrogenase | 0.012 | 2.167 | -0.177 | 0.842 |
| sco4139 | Phosphate transporter ATP-binding protein | -0.530 | 2.0167 | 0.811 | -0.115 |
| sco4140 | Phosphate ABC transporter permease | -0.212 | 2.510 | 0.440 | -0.180 |
| sco4141 | Phosphate ABC transporter permease | 0.0242 | 2.199 | 0.715 | -0.039 |
| sco4142 | Phosphate-binding protein | -0.445 | 2.474 | 0.741 | 0.128 |
| sco7023 | Hypothetical protein | -0.126 | 2.033 | -0.096 | 0.454 |
| sco5265 | Hypothetical protein | 0.056 | -2.274 | -0.029 | 0.056 |
| sco5338 | Regulatory protein | -0.723 | -3.560 | 0.195 | 0.0240 |
| sco5339 | Plasmid transfer protein | -0.486 | -2.302 | 0.476 | -0.078 |
| sco0185 | Geranylgeranyl pyrophosphate synthase | 0.685 | 0.555 | -0.500 | 0.273 |
| sco0186 | Phytoene dehydrogenase | 0.322 | 0.329 | 0.802 | 0.557 |
| sco0187 | Phytoene synthase | 0.580 | 0.431 | -2.672 | -0.315 |
| sco0191 | Lycopene cyclase | 0.0560 | 0.424 | -2.626 | 0.039 |
| sco0192 | Oxidoreductase | 0.194 | 0.459 | -2.232 | 0.232 |
| sco0193 | DNA-binding regulator | 0.303 | 1.144 | -2.137 | 1.342 |
| sco3706 | ABC transporter ATP-binding protein | -0.117 | 0.355 | -0.2690 | 2.149 |
| sco3705 | ABC transporter membrane subunit | 0.039 | 0.431 | 0.218 | 2.461 |
| sco3704 | Substrate-binding transport protein | -0.203 | 0.140 | 0.0916 | 2.229 |

* Differentially expressed genes at least one condition between the Δ*rokL6* and wt; sample collected from MM+mannitol at 24 h (VEG-Mann), MM+mannitol at 48 h (SPO-Mann), MM+mannitol and GlcN at 24 h (VEG-GlcN), MM+mannitol and GlcN at 48 h (SPO-GlcN).

**Table S2.** Putative targets of RokL6 as predicted by PREDetector

| Gene | Annotation | Scores # | Sequence* |
|---|---|---|---|
| sco1448 | Probable transport protein | 29.4 | CTATCAGGCAGGCTCCCTGATAG |
| *rokL6* | ROK family regulator | 29.4 | CTATCAGGCAGGCTCCCTGATAG |
| sco0137 | Transmembrane transport protein | 15.4 | CTTTCAGACATGGTTCCTGATTG |
| sco4114 | Sporulation associated protein | 7.1 | TTAAGAGGCATTGTTCCGGATGG |
| sco1359 | Possible integral membrane protein | 6.7 | CTACGAGGACGACTGCCTCATCG |
| sco1358 | LysR family transcriptional regulator | 6.7 | CTACGAGGACGACTGCCTCATCG |
| sco5864 | Hypothetical protein | 6.2 | TGATCATGCACCCTGTCGGAAAG |
| sco3250 | Probable integrase | 6.0 | CCATCAGGCAGCCTCCTCGATCT |

* Sections of sequences underlined are those that match the proposed RokL6 consensus sequence.
### Matrix scores determined by PREDetector; score above 10 are considered relevant

#### Supplementary Figures

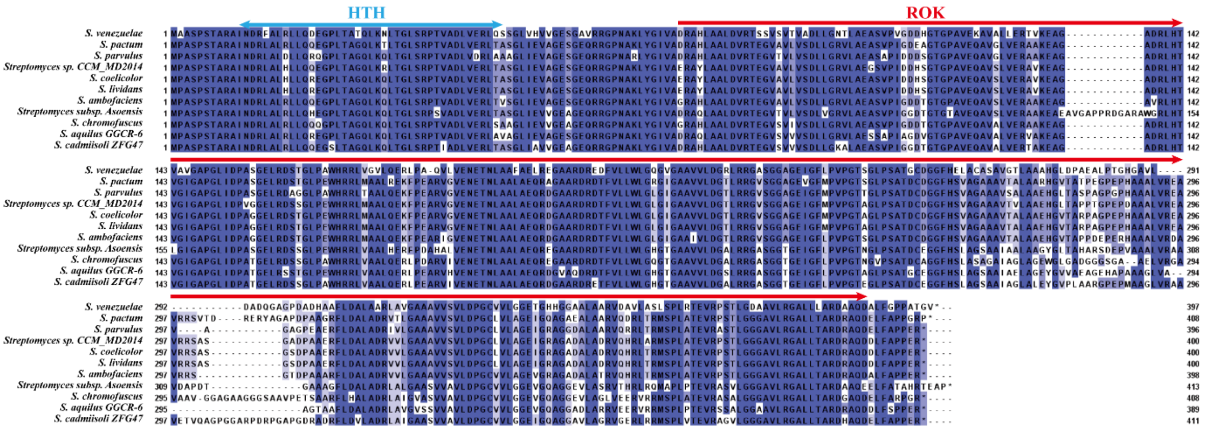

**Fig. S1. Alignment of RokL6 and orthologs in *Streptomyces*.** Multiple sequence alignment was performed using the amino acid sequence of RokL6 and its orthologs from other *Streptomyces* species by Clustal Omega. Identical amino acids are shown in dark blue, and amino acids with similar properties in light blue. The helix-turn-helix (HTH) domain and ROK-kinase domain are indicated. The image was generated using Jalview (version 2.11.2.7).

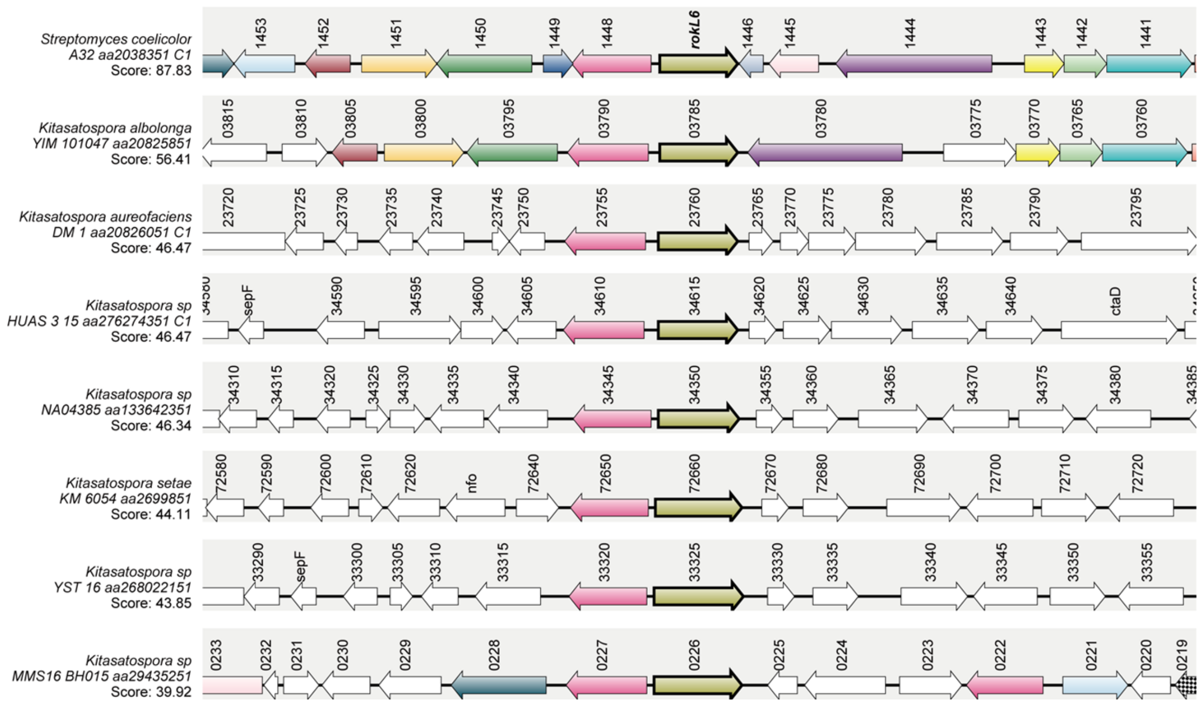

**Fig. S2. Gene synteny analysis of *rokL6*-sco1448 in *Kitasatospora* species**. Synteny analysis was performed by Synttax (scores are given). Homologous genes are presented as indicated and in the same colours.

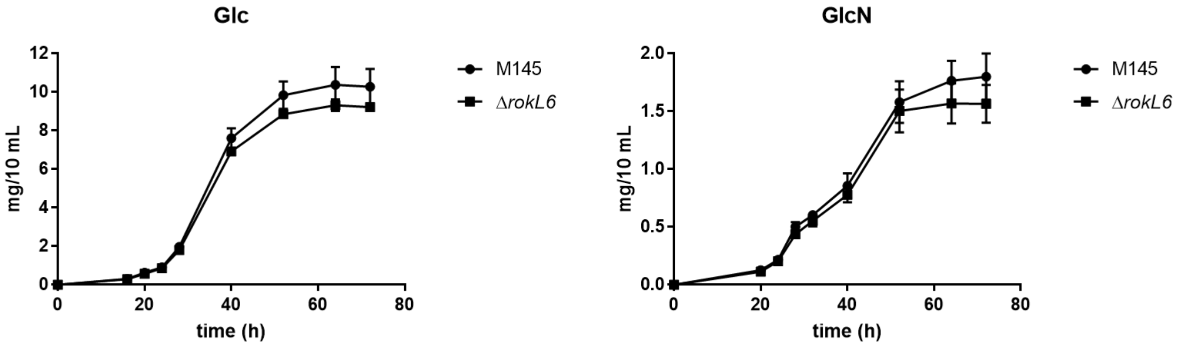

**Fig. S3. Growth curves of M145 and Δ*rokL6* in NMMP.** Comparison of the growth pattern between wild-type *S. coelicolor* M145 and the mutant Δ*rokL6*. Strains were growth in liquid NMMP supplemented with either 1% (w/v) glucose (Glc) or glucosamine (GlcN). The dry weights of the mycelia were measured at different time points. Each value was obtained from three biological replicates as a mean ± SD.

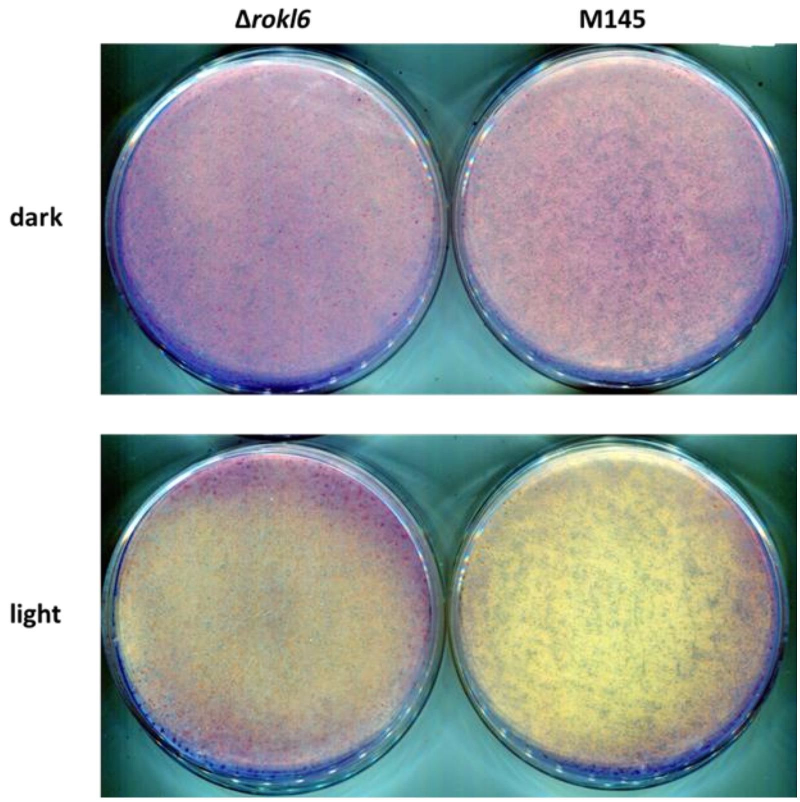

**Fig. S4. RokL6 up-regulates production of carotenoids.** $10^7$ spores of wild-type M145 and its *rokL6* mutant were plated on MM with 1% mannitol and 50 mM GlcN, and cultured at 30˚C for 48 h in the dark (dark) and light (light). Note that the petri dish plated with wild-type M145 appeared significantly yellower than that with Δ*rokL6* due to the upregulated production of carotenoid under the light condition (no significant difference observed in the dark).

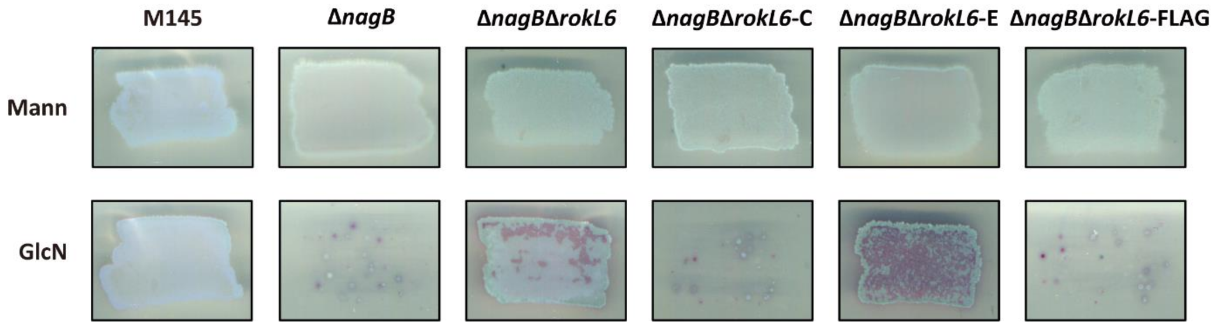

**Fig. S5. Verification of the functionalily of RokL6-FLAG$_3$.** Spores of *S. coelicolor* M145, its mutants Δ*nagB*, Δ*nagB*Δ*rokL6*, Δ*nagB*Δ*rokL6*-C (Δ*nagB*Δ*rokL6* complemented with RokL6), Δ*nagB*Δ*rokL6*-FLAG (Δ*nagB*Δ*rokL6* complemented with RokL6-FLAG$_3$) and Δ*nagB*Δ*rokL6* harbouring empty plasmid pSET152 (Δ*nagB*Δ*rokL6*-E) were plated on MM with either mannitol (Mann) or mannitol with GlcN (GlcN), and cultured at 30˚C to check forr growth on GlcN. Note that introduction of a plasmid expressing either wild-type RokL6 (column 4) or RokL6-FLAG$_3$ (column 6) made the Δ*nagB*Δ*rokL6* double mutant sensitive to GlcN once more.

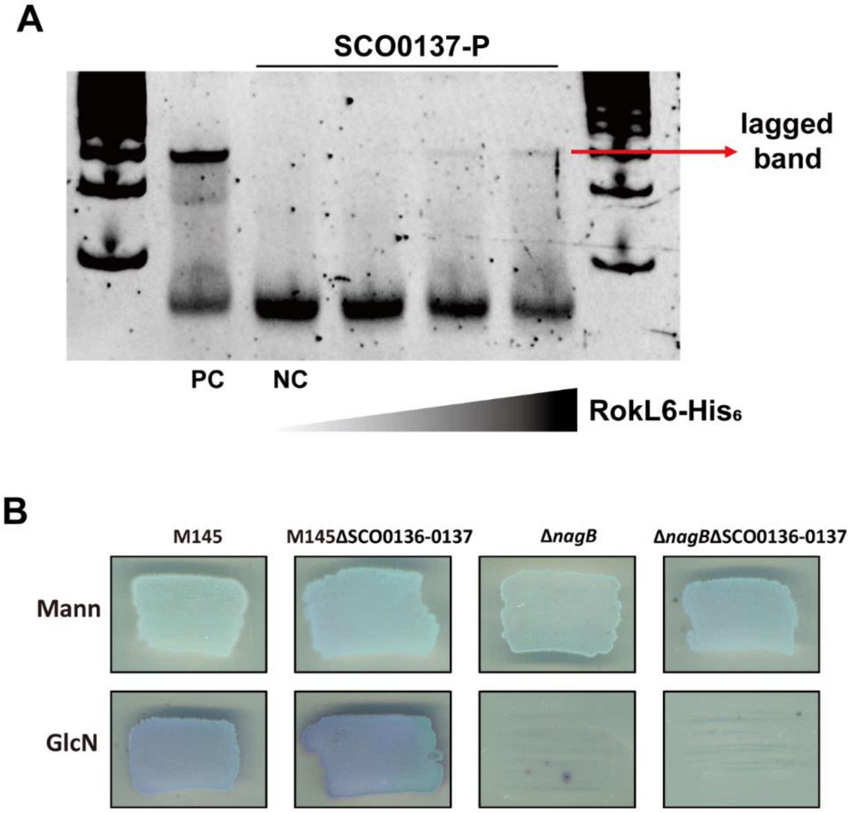

**Fig. S6. Binding of RokL6 to the sco0137 promoter region sco0137-P.** (A) EMSA of RokL6-His$_6$ with the upstream region of the sco0137-0136 operon. The *rokL6* promoter region with 0.5 µM RokL6-His$_6$ was used as positive control (PC), and sco0137-P without RokL6-His$_6$ was used as negative control (NC); lagged band is indicated by red arrow. Note that binding signal of RokL6 to the promoter is very weak; (B) Growth of sco0136-0137 mutants on GlcN. *S. coelicolor* M145, Δsco0136-0137, Δ*nagB* and Δ*nagB*Δsco0136-0137, were grown on MM with mannitol (Mann), and MM with mann and GlcN (GlcN) for their viability evaluation. Note that deletion of sco0136-0137 in Δ*nagB* did not relieve the toxicity of GlcN.

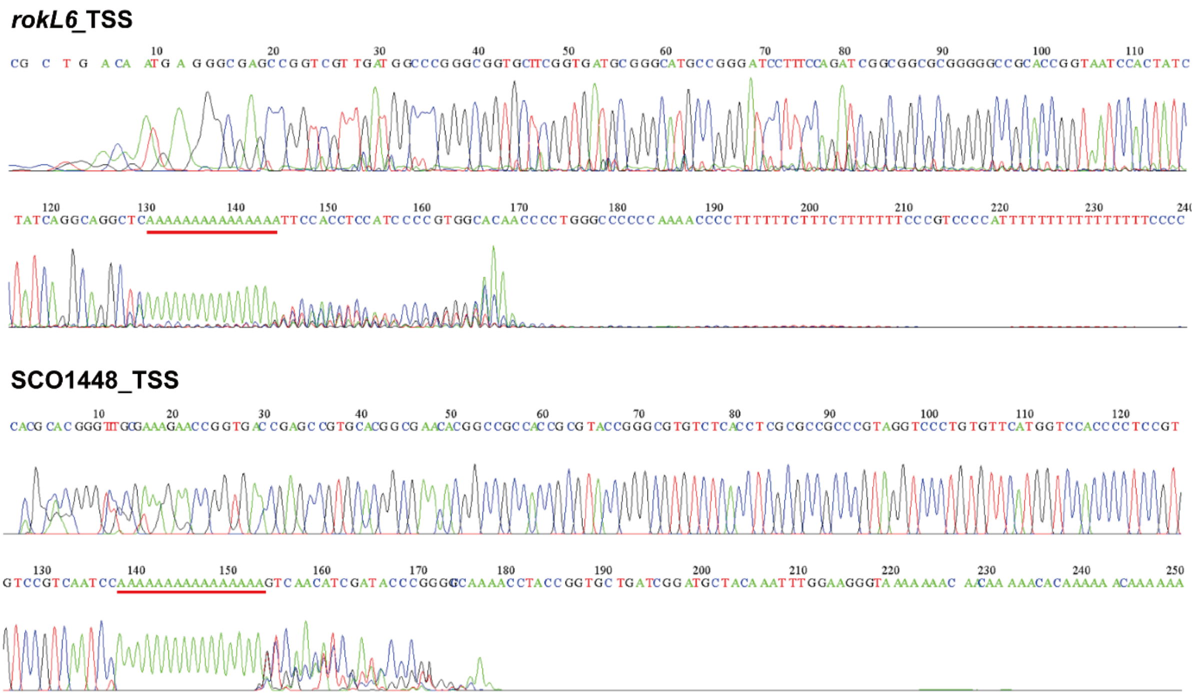

**Fig. S7. Determination of TSSs of *rokL6* and sco1448 by 5' RACE.** Red lines: complementary sequences of oligo(dT) anchor primer.
